# The Arf-GAP Age2 localizes to the late-Golgi via a conserved amphipathic helix

**DOI:** 10.1101/2023.07.23.550229

**Authors:** Kaitlyn M. Manzer, J. Christopher Fromme

## Abstract

Arf GTPases are central regulators of the Golgi complex, which serves as the nexus of membrane trafficking pathways in eukaryotic cells. Arf proteins recruit dozens of effectors to modify membranes, sort cargos, and create and tether transport vesicles, and are therefore essential for orchestrating Golgi trafficking. The regulation of Arf activity is controlled by the action of Arf-GEFs, which activate via nucleotide exchange, and Arf-GAPs, which inactivate via nucleotide hydrolysis. The localization dynamics of Arf GTPases and their Arf-GAPs during Golgi maturation have not been reported. Here we use the budding yeast model to examine the temporal localization of the Golgi Arf-GAPs. We also determine the mechanisms used by the Arf-GAP Age2 to localize to the Golgi. We find that the catalytic activity of Age2 and a conserved sequence in the unstructured C-terminal domain of Age2 are both required for Golgi localization. This sequence is predicted to form an amphipathic helix and mediates direct binding of Age2 to membranes *in vitro*. We also report the development of a probe for sensing active Arf1 in living cells and use this probe to characterize the temporal dynamics of Arf1 during Golgi maturation.

## Introduction

The Golgi complex plays a central role in both the secretory and endocytic pathways of eukaryotic cells by directing the proper sorting, modification, and transport of proteins to their appropriate cellular destinations. To achieve this, the Golgi complex facilitates several incoming and outgoing trafficking routes that occur at specific stages throughout cisternal maturation to ensure the timely and accurate delivery of cargo (Pantazopoulou & Glick, 2019).

A key player in this process is the small GTPase Arf1, which functions as a molecular switch, transitioning between an inactive GDP-bound state and an active GTP-bound state (Jackson & Bouvet, 2014). Inactive Arf1 is cytosolic, and once active, Arf1 undergoes a conformational change that stabilizes it on the Golgi membrane and allows it to recruit various effectors. In budding yeast, Arf1 has a close paralog, Arf2, that is 96% identical, genetically redundant, and expressed at ∼10% the level of Arf1 (Stearns et al., 1990). In humans, Arf1 has three paralogs (Arf3, Arf4, and Arf5) that have apparently distinct but genetically redundant roles in Golgi trafficking (Pennauer et al., 2022; Volpicelli-Daley et al., 2005; Wong-Dilworth et al., 2023).

Arf1 is responsible for recruitment of the machinery needed to form vesicles destined for the lysosome (AP-3) and endosomes (AP-1, GGA), as well as recycling back to earlier Golgi compartments or the ER (COPI, AP-1) (Adarska et al., 2021; Casler et al., 2019; Dell’Angelica et al., 2000; Donaldson et al., 1992; Ooi et al., 1998; Stamnes & Rothman, 1993; Traub et al., 1993). In the budding yeast *Saccharomyces cerevisiae*, Arf1 also recruits exomer for the transport of cargo to the plasma membrane (PM), although no homolog of exomer has been identified in metazoans (Paczkowski et al., 2015; Sanchatjate & Schekman, 2006; Trautwein et al., 2006; Wang et al., 2006). Given its central role in initiating several different trafficking events at the Golgi, it is not surprising that Arf1 is also important for the overall Golgi maturation process (Bhave et al., 2014).

To ensure that these vesicles are formed at the proper time and sort the correct cargos, Arf1 is tightly regulated. The state of Arf1 is dictated by guanine nucleotide exchange factors (GEFs) and GTPase-activating proteins (GAPs) (Casanova, 2007; Spang et al., 2010). Arf GTPases possess very low intrinsic rates for GTP to GDP exchange and have undetectable GTPase activity on their own (East & Kahn, 2011). GEFs activate GTPases by facilitating the exchange of bound GDP for GTP, while GAPs inactivate GTPases by stimulating GTP hydrolysis to generate GDP. As active Arf1 binds cargo adaptors and coat proteins, the inactivation of Arf1 by Arf-GAPs is thought to be required for the release of the coat and subsequent fusion of the vesicle with its target membrane. Arf-GAPs have also been implicated in cargo selection and coat assembly (Y. Shiba & Randazzo, 2012).

The Arf-GEFs and Arf-GAPs reside on specific membranes, allowing for tight spatial control of Arf1. The differential localization of these regulatory proteins is a key aspect of Arf1 regulation. In yeast, there are three Arf1-GEFs that localize to the Golgi. Gea1 and Gea2 (paralogous to human GBF1) reside at the early- and medial-Golgi respectively to activate Arf1 for intra-Golgi and Golgi-to-Endoplasmic Reticulum (ER) retrograde transport (Gustafson & Fromme, 2017; Spang et al., 2001). Sec7 (paralogous to human BIG1/2) localizes to the late-Golgi to activate Arf1 for secretion and transport to endolysosomal compartments (Bui et al., 2009). These Arf-GEFs are thought to localize to their respective locations via recruitment by GTPases and preference for membranes of specific character (Gustafson & Fromme, 2017; McDonold & Fromme, 2014).

Arf1 is inactivated by four Arf-GAPs in yeast: Gcs1, Glo3, Age2, and Age1, which are paralogous to the human ARFGAP1, ARFGAP2/3, SMAP2, and ACAP1 proteins, respectively. Gcs1 harbors an “ALPS” motif that preferentially binds curved membranes for localization (Bigay et al., 2005; Drin et al., 2007; Mesmin et al., 2007), and Glo3 requires an interaction with COPI for its localization (Eugster et al., 2000; Kliouchnikov et al., 2009; Schindler et al., 2009; Xie et al., 2023). However, the temporal localization dynamics of the Arf-GAPs during Golgi maturation has not been established.

Furthermore, the localization dynamics of active Arf1 at the Golgi remains unclear. Fusion of fluorescent proteins to Arf1 impairs the function of Arf1 (Jian et al., 2010), and thus there is currently no method for the visualization of functional Arf1 in live cells during the Golgi maturation process in yeast. Ascertaining the dynamics of the localization of Arf1 and its regulators is key to our understanding of Arf1-mediated trafficking.

Here, we provide a spatiotemporal analysis of the Arf-GAPs during Golgi maturation in *S. cerevisiae*, in which Glo3, Gcs1, and Age2 were found to mainly localize at the early-, medial-, and late-Golgi respectively. We also report a previously unknown mechanism for the localization of the late-Golgi Arf1-GAP, Age2. Surprisingly, despite a reported interaction of SMAP2 with clathrin (Natsume et al., 2006), neither clathrin nor its adaptors were required for Age2 localization to the Golgi. Instead, we identified a highly conserved amphipathic helix in Age2 that contributes to its targeting through direct interaction with the membrane. Interestingly, the ability of Age2 to act on its substrate, Arf1, was also found to play a role in achieving Age2 localization. This feature appears unique to Age2, as Gcs1 and Glo3 do not seem to require an interaction with Arf1 for its localization. Finally, we developed novel methods for the direct visualization of Arf1 in living cells without disrupting Arf1 function. Using these approaches, we determined that active Arf1 is enriched at the late Golgi, likely to support the high demand for Arf1 in several outgoing transport pathways at the *trans*-Golgi network (TGN).

## Results

### An Arf-GAP is present at each stage of Golgi maturation

The four Arf1 GAPs in *S. cerevisiae*, Gcs1, Glo3, Age2, and Age1 (Poon et al., 1996, 1999, 2001; Zhang et al., 2003), each possess a conserved catalytic GAP domain, with the rest of their domain structures varying significantly (Spang et al., 2010), allowing for functional specificity (Figure 1A). Although the approximate localization of these Arf-GAPs has been reported (Benjamin et al., 2011; Funaki et al., 2011; Liu et al., 2005; Natsume et al., 2006; M. Robinson et al., 2006; Weimer et al., 2008), to better understand their dynamics, we sought to investigate their temporal localization throughout Golgi maturation.

**FIGURE 1.**
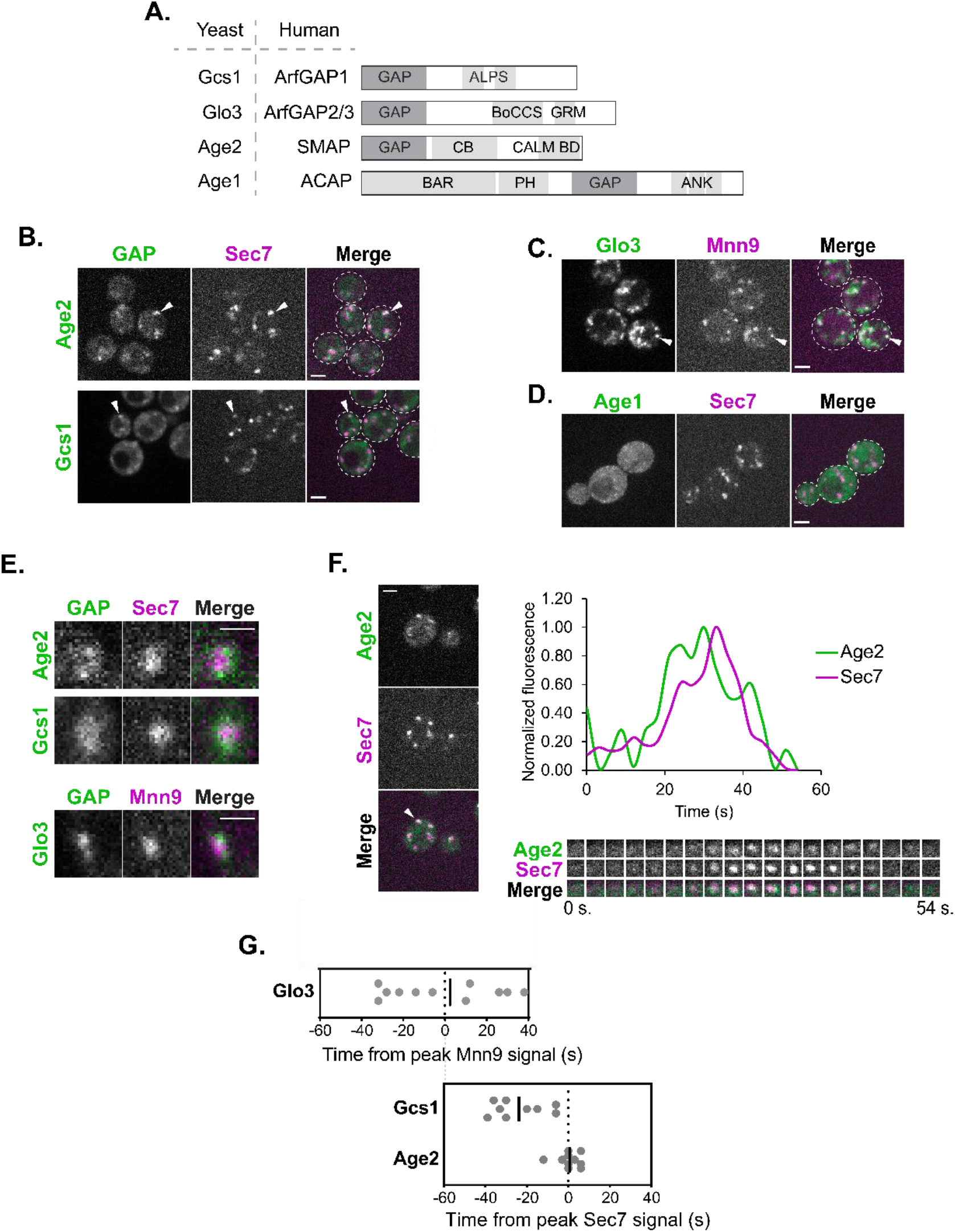
The Arf-GAPs Glo3, Gcs1, and Age2 localize at the early-, medial-, and late-Golgi respectively. (A) Schematic of the yeast Arf-GAPs and their human homologs and domain structures. GAP, GAP domain; ALPS, Amphipathic Lipid Packing Sensor; BoCCS, Binding of Coatomer, Cargo and SNARE; GRM, Glo3 Regulatory Motif; CB, Clathrin Binding; CALM BD, CALM Binding Domain; BAR, Bin/Amphiphysin/Rvs; PH, Pleckstrin Homology Domain; ANK, Ankyrin Repeat. (B) Subcellular localization of Age2-GFP or Gcs1-GFP relative to the late-Golgi marker Sec7-6xDsRed. Arrowheads indicate colocalization between tagged proteins. Single focal plane. (C) Subcellular localization of Glo3-Neon relative to the early-Golgi marker Mnn9-mCherry. Arrowheads indicate colocalization between tagged proteins. Single focal plane. (D) Subcellular localization of endogenous Neon-tagged Age1. Maximum projection. (E) Magnified images showing the distribution of endogenously tagged GAPs (Age2-GFP, Gcs1-Neon, and Glo3-Neon) at single Golgi compartments (Mnn9-mCherry, early-Golgi, or Sec7-6xDsRed, late-Golgi). Scale bar represents 1 µm. (F) Left: Representative image of time-lapse microscopy of Age2-GFP versus Sec7-6xDsRed. Arrowhead denotes Golgi compartment of interest. Single focal plane. Bottom right: Imaging of the compartment of interest over time. Top right: Plot of normalized fluorescence intensity in the compartment of interest over time. (G) (Top) Peak-to-peak times (s) for Glo3 versus Mnn9 and (Bottom) for Gcs1 and Age2 versus Sec7. Line represents the mean. Offset between two graphs is for visual purposes, and was approximated based on time-lapse analyses reported in Sardana et al., 2021 and Tojima et al., 2019. For all images, scale bars represent 2 µm unless otherwise noted.

We began by C-terminally tagging the Arf-GAPs on the chromosome at their endogenous locus. We verified that tagged Gcs1, Glo3, and Age2 each retained their essential function, as indicated by the viability of tagged strains in sensitized backgrounds that would be inviable if Arf-GAP function were lost (Supplemental Figure S1A). We were unable to perform a similar functional analysis with Age1 as the *AGE1* gene has no known synthetic lethal interactions.

To determine which Golgi proteins would be most suitable for comparative time-lapse analysis for each Arf-GAP, we confirmed the localization of the Arf-GAPs relative to established marker proteins. As expected, we observed several instances of colocalization of Gcs1 and Age2 with the late-Golgi marker, Sec7 (Figure 1B) (Losev et al., 2006). Glo3 instead colocalized well with the early-Golgi marker, Mnn9 (Figure 1C), as well as the COPI α subunit (yeast Cop1 protein) (Supplemental Figure S1B), consistent with its known role in COPI function (Poon et al., 1999; Weimer et al., 2008). Age1 exhibited only faint and subtle localization to very small punctate structures, likely due to its low expression level (Ho et al., 2018), and therefore we were unable to analyze endogenously expressed Age1 (Figure 1D). We thus proceeded with time-lapse analysis of Gcs1 and Age2 relative to Sec7, and Glo3 relative to Mnn9. We observed that each of the three Arf-GAPs tended to localize towards the periphery of the Golgi compartments, often as multiple small punctate dots, perhaps representing sites of vesicle formation (Figure 1E).

We measured the fluorescence intensity over time of each Arf-GAP relative to that of the chosen marker at individual Golgi compartments (Losev et al., 2006; Matsuura-Tokita et al., 2006). Age2 signal peaked at approximately the same time as Sec7 (Figure 1, F and G). Several clathrin-mediated trafficking pathways occur at the late-Golgi, and thus this is consistent with the role of Age2 in clathrin-mediated post-Golgi transport (Poon et al., 2001).

Peak Gcs1 localization occurred upstream of Sec7 and became undetectable before the end of the Sec7 compartment’s lifetime suggesting that Gcs1 predominantly functions at the medial-Golgi (Figure 1G, Supplemental Figure S1C). We observed instances of colocalization of Gcs1 with markers of the early-, medial, and late-golgi. This is consistent with the reported function of Gcs1 in both early-Golgi-to-ER trafficking as well as post-Golgi trafficking (Poon et al., 1999, 2001).

Glo3 was found to localize broadly throughout the earlier stages of Golgi maturation (Supplemental Figure S1D; Figure 1G). This is in agreement with previous studies that found that Glo3 directly interacts with COPI (Eugster et al., 2000; Lewis et al., 2004; Schindler et al., 2009; Xie et al., 2021, 2023), and that COPI vesicles form for both intra-Golgi and Golgi-to-ER transport earlier in Golgi maturation (Papanikou et al., 2015; Tojima et al., 2019). These time-lapse analysis findings were further corroborated by conventional colocalization analysis (Supplemental Figure S2, A and B).

To gain insight into where Age1 may be functioning, we overexpressed tagged Age1 on a plasmid under the strong ADH1 promoter. While still dim, we were able to observe punctate structures that overlapped with Sec7 compartments but not Mnn9 compartments, suggesting that Age1 likely also functions at the late Golgi (Supplemental Figure S2C), although its low expression level and lack of synthetic lethal interactions indicates it is unlikely to play an important role during vegetative growth under standard laboratory conditions.

Of note, the maturation dynamics of Gcs1 and Glo3 are also consistent with the reported localizations of the human homologs ArfGAP1 and ArfGAP2/3. An ultrastructural localization analysis reported that ArfGAP1 was found distributed equally between the *cis-* and *trans*-Golgi, while ArfGAP2/3 had a greater distribution at the *cis*-Golgi (Weimer et al., 2008). Therefore, localization is a conserved feature of these Arf-GAPs.

### Age2 localization and function requires the GAP domain and the KKSILSLY motif

The differential localization of the Arf-GAPs likely reflects an important aspect of Arf1 regulation. Therefore, we sought to understand the mechanism for the localization of each of the Arf-GAPs to their respective membranes. Major determinants for targeting have already been identified for both Glo3 and Gcs1 (Arakel et al., 2019; Spang et al., 2010; Weimer et al., 2008). Glo3 requires an interaction with COPI for localization (Eugster et al., 2000; Kliouchnikov et al., 2009), and Gcs1 requires an interaction with the membrane through a membrane curvature-sensing motif termed the ALPS motif (Bigay et al., 2005; Drin et al., 2007).

In contrast, the localization mechanism for Age2 is largely unknown. The human homolog SMAP2 has been reported to possess a clathrin binding (CB) domain and a CALM binding domain (Natsume et al., 2006). The CB domain of SMAP2 harbors two clathrin interacting motifs, a canonical clathrin box motif and an atypical DLL motif (Natsume et al. 2006). The DLL motif is absent in Age2, and the sequence corresponding to the clathrin box motif in Age2 does not match the canonical clathrin box motif (LΦXΦ[DE], where x is any amino acid and Φ is a bulky hydrophobic residue) (Supplemental Figure S3A) (Smith et al., 2017). However, variants of the clathrin box motif have been identified that retain the ability to bind clathrin (Dell’Angelica, 2001), and thus we consider it possible that Age2 also binds clathrin. The C-terminal portion of SMAP2 that is responsible for the CALM interaction is not conserved in Age2 (Supplemental Figure S3A).

To gain insight into which regions of Age2 may be important for its function and localization, we examined sequence alignments of Age2 with its homologs in other species (Figure 2A). We identified a highly conserved “KKSILSLY” sequence within the region of the protein that is C-terminal to the GAP domain. To test if this sequence plays a role in Age2 localization, we created a mutant in which the KKSILSLY sequence was replaced with alanine residues (KKSILSLY mutant). This resulted in a complete redistribution of Age2 to the cytosol (Figure 2B). We verified that this mutant was expressed, and thus this effect was not due to degradation or a lack of expression (Supplemental Figure S4A). Therefore, the KKSILSLY sequence is required for Age2 localization to the Golgi.

**FIGURE 2.**
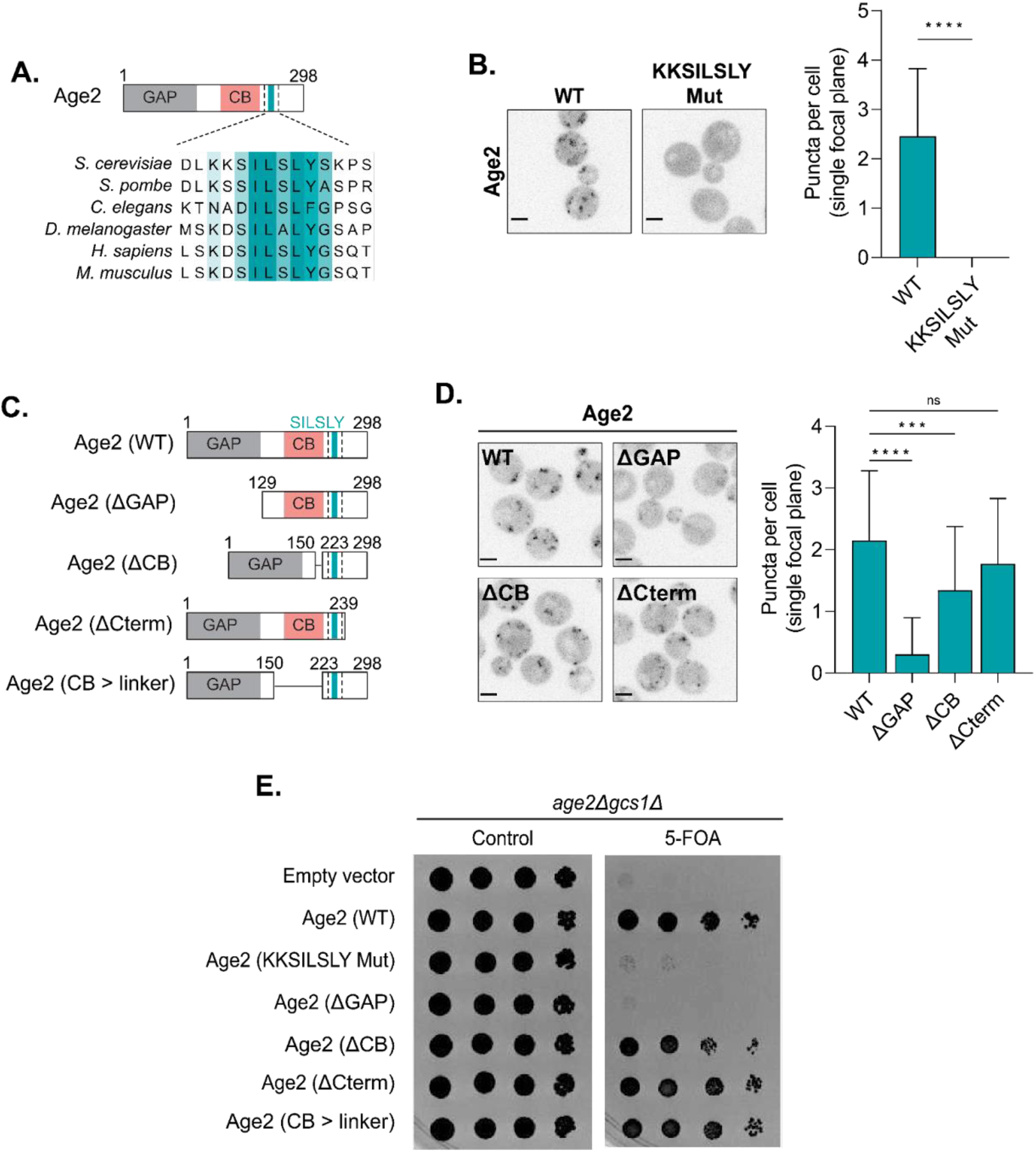
Dissection of Age2 functional domains. (A) Alignment highlighting conserved KKSILSLY sequence. GAP, GAP domain; CB, Clathrin Binding. (B) Left: Fluorescence microscopy of WT Age2-Neon-3xFLAG and Age2 (KKSILSLY mutant)-Neon-3xFLAG. Maximum projections. Right: Quantification of the number of puncta in a single focal plane per cell. Error bars represent standard deviation for n > 22 cells quantified. (C) Diagram depicting Age2 truncations. GAP, GAP domain; CB, Clathrin Binding. (D) Left: Fluorescence microscopy of Age2-Neon-3xFLAG truncations. Maximum projections. Right: Quantification of the number of puncta in a single focal plane per cell. Error bars represent standard deviation for n > 22 cells quantified. (E) Complementation assay with Age2-Neon-3xFLAG mutants and truncations. Scale bars represent 2 µm. n.s., not significant; *, P ≤ 0.05, **, P ≤ 0.01, ***, P < 0.001; ****, P < 0.0001.

To determine if the KKSILSLY sequence might be sufficient for protein localization or if there are other regions of Age2 that are also required for its localization, we performed a truncation analysis. We designed several truncation constructs, in which we removed either the GAP domain (Age2 ΔGAP), the central portion of the protein that harbors the putative clathrin box motif (Age2 ΔCB), or the C-terminus downstream of the KKSILSLY sequence (Age2 ΔCterm) (Figure 2C).

We found that removal of the GAP domain resulted in Age2 largely redistributing to the cytosol, indicating that the GAP domain itself is important for Age2 localization (Figure 2D). We verified that the Age2 ΔGAP mutant was still expressed (Supplemental Figure S4A).

Removal of the C-terminal ∼50 residues had no significant impact on Age2 localization (Figure 2D). In contrast, we observed that loss of the CB domain resulted in a subtle, but significant perturbation of Age2 localization (Figure 2D). This suggests that this region plays a minor role in Age2 localization to the Golgi.

To assess which portions of the protein are required for Age2 function, we performed a complementation assay using sensitized *gcs1Δ* cells (Figure 2E). Age2 is not essential in otherwise wild-type cells, but the combined loss of Age2 and Gcs1 results in synthetic lethality (Poon et al., 2001). As expected, loss of the GAP domain, which is essential for the GAP activity of Age2, resulted in a lack of growth. Mutation of the KKSILSLY sequence also resulted in a complete lack of growth. Removal of the CB resulted in a minor growth defect, but growth was rescued by the substitution of this region with a flexible linker (Age2 CB > linker), suggesting the importance of the CB for cell growth in this background is in its length (Figure 2, C and E). The loss of the C-terminal region downstream of the KKSILSLY sequence was dispensable for cell growth (Figure 2E). Taken together these results establish the requirement of the GAP domain and KKSILSLY sequence for proper localization and function of Age2.

### The KKSILSLY sequence forms a membrane-binding amphipathic helix

To further define the role of the KKSILSLY sequence in Age2 localization, we first substituted the individual residues within the sequence to alanine (Figure 3A). We found that substitution of both lysine residues or of any hydrophobic residue within the sequence resulted in significant effects on localization. Several Arf-GAPs and Arf-GEFs possess membrane-targeting amphipathic helices (AH) (Bigay et al., 2005; Drin et al., 2007; Kliouchnikov et al., 2009; Muccini et al., 2022). When we further examined the KKSILSLY sequence, it did appear likely to fold into an amphipathic helix based on helical wheel analysis (Figure 3B) and AlphaFold (Jumper et al., 2021; Varadi et al., 2021) structure prediction (Supplemental Figure S4B). Note that lysine residues are known to “snorkel”, with the charged side chain associating with phospholipid head groups (Drin & Antonny, 2010).

**FIGURE 3.**
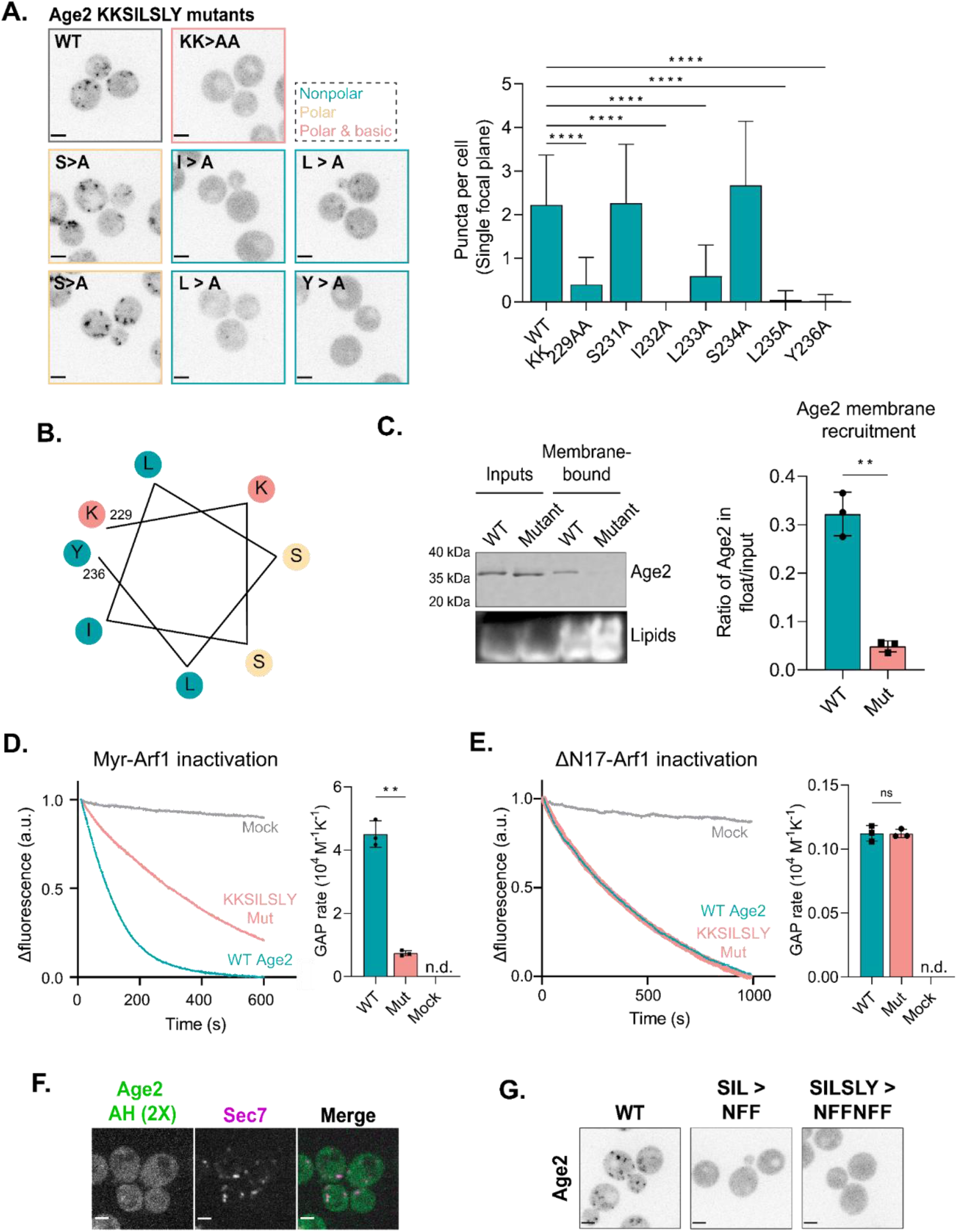
The KKSILSLY sequence in Age2 forms an amphipathic helix that mediates membrane-binding to inactivate Arf1. (A) Left: Fluorescence microscopy of Neon-3xFLAG-tagged Age2 point mutants. Maximum projections. Border colors correspond to the properties of the mutated residues, where teal is nonpolar, yellow is polar and neutral, and pink is polar and basic. Right: Quantification of the number of puncta in a single focal plane per cell. Error bars represent standard deviation for n > 22 cells quantified, n.s. not shown. (B) Helical wheel of the KKSILSLY sequence (residues 229 -236). The properties of each amino acid are represented by color, where teal is nonpolar, yellow is polar and neutral, and pink is polar and basic. (C) Left: Representative *in vitro* liposome flotation assay to assess binding of purified wild-type and KKSILSLY mutant Age2 to membranes. Right: Quantification of Age2 membrane recruitment, as measured by coomassie band intensity ratio, for n = 3 assays. Error bars represent standard deviation. (D) Left: *In vitro* GAP assay fluorescence traces for wild-type and KKSILSLY mutant Age2 GAP activity on myr-Arf1 bound to liposome membranes. A.u, arbitrary units. Right: Quantification of GAP activity rates. Error bars represent standard deviation. N.d., not detected. (E) *In vitro* GAP assay of wild-type and KKSILSLY mutant Age2 GAP activity on ΔN17-Arf1 in solution (without liposomes). Error bars represent standard deviation. N.d., not detected. A.u, arbitrary units. (F) Fluorescence microscopy of tandem repeats of Age2 AH (a portion of Age2 including the KKSILSLY motif) tagged with Neon expressed on a centromeric plasmid alongside Sec7-6xDsRed. Single focal plane shown. (G) Fluorescence microscopy of Age2-Neon-3xFLAG mutants, in which mutations were introduced to alter the sequence while maintaining its amphipathic character. Maximum projections. Scale bars represent 2 µm. n.s., not significant; *, P ≤ 0.05, **, P ≤ 0.01, ***, P < 0.001; ****, P < 0.0001.

To directly test if this sequence is used by Age2 to bind the membrane surface, we purified both wild-type and KKSILSLY mutant Age2 constructs and assessed their ability to bind synthetic liposome membranes *in vitro*. Age2 exhibited an affinity for membranes that was significantly reduced by the KKSILSLY mutation, indicating a direct role of this sequence in Age2 membrane binding (Figure 3C).

We next asked whether the membrane association conferred by this amphipathic helix is required for Arf-GAP activity towards membrane-bound Arf1-GTP. We performed *in vitro* GAP assays using previously established methods (Antonny, Huber, et al., 1997; Paczkowski et al., 2012; Richardson & Fromme, 2015) to assess the GAP activity of Age2 on active myristoylated Arf1 (myrArf1) bound to liposomes. This assay involves measuring the intrinsic tryptophan fluorescence of Arf1 to monitor changes in its conformational state. We observed that the KKSILSLY mutant exhibited a significantly reduced ability to inactivate Arf1, with a more than 6-fold reduction in GAP activity compared to the wild-type (Figure 3D; Supplemental Figure S4C). Notably, the KKSILSLY mutant retained the ability to inactivate soluble Arf1 (ΔN17-Arf1) in the absence of membranes, exhibiting a similar rate of GAP activity to that of the wild-type under these conditions (Figure 3E, Supplemental Figure S4D). Taken together, these results indicate that the KKSILSLY sequence of Age2 forms a conserved amphipathic helix that facilitates binding of Age2 to the Golgi membrane in order to inactivate Arf1.

The *in vivo* truncation results suggested that this amphipathic helix is essential but not sufficient for localization. We postulated that expressing two repeats of a short sequence that includes the KKSILSLY sequence in tandem might increase its affinity for the Golgi membrane enough to obviate the requirement of the GAP domain for Age2 localization. While the tandem AH construct (Age2 AH 2X) did exhibit some punctate localization that overlapped with the late-Golgi, the puncta were rare and faint (Figure 3F). This result further underscores the importance of the GAP domain in Age2 localization.

Lastly, we tested if the precise primary sequence of the amphipathic helix was important, or if amphipathicity provided by different residues might be sufficient for membrane localization. We created a mutant in which KKSILSLY was replaced with KKNFFNFF, altering the sequence while maintaining its amphipathic character. This substitution completely abolished the Golgi localization of Age2 (Figure 3G). Therefore, the primary sequence requirements of this motif are specific, explaining the strong conservation of this sequence across species.

### GAP activity is important for Age2 localization

We next investigated the observation that a loss of the catalytic GAP domain impaired Age2 localization (Figure 2D). We wondered if this effect was due to a loss of GAP activity or if the GAP domain participates in additional interactions that contribute to its recruitment. We therefore mutated the catalytic Arg residue (R52K), rendering Age2 ‘GAP-dead’. Interestingly, the GAP-dead Age2 R52K mutant was significantly mislocalized despite being expressed normally (Figure 4A, Supplemental Figure S4A). The same result was observed when Age2 R52K was expressed as an extra copy in cells also expressing wild-type Age2 (Supplemental Figure S4E), and thus mislocalization does not appear to be due to a downstream effect of loss of Age2 function in cells. Mislocalization was also observed if the Arg residue was mutated to alanine (Age2 R52A) (Supplemental Figure S4F).

**FIGURE 4.**
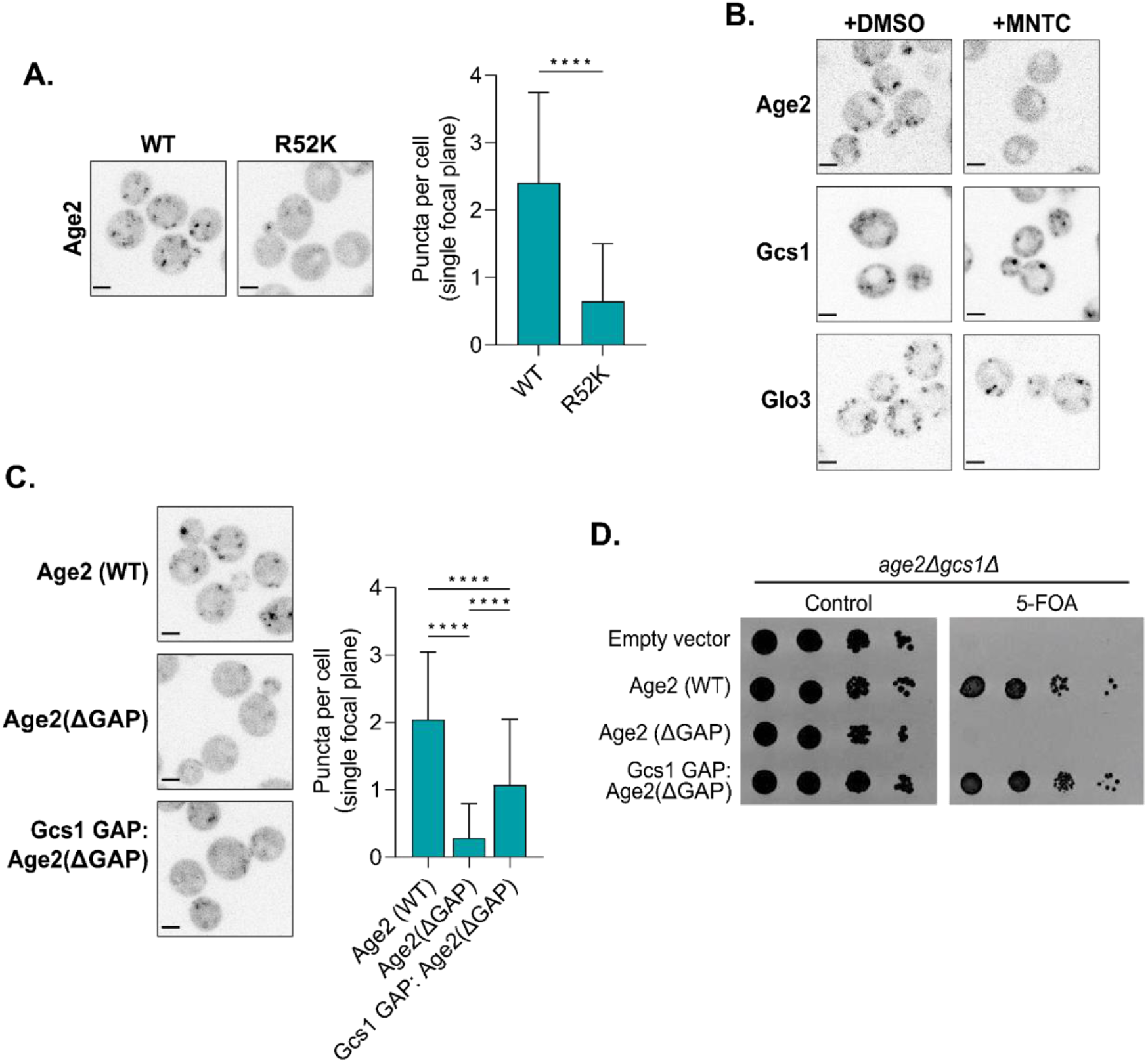
The interaction of Age2 with Arf1 is important for Age2 localization. (A) Left: Fluorescence microscopy of GAP dead Age2 (R52K)-Neon-3xFLAG. Maximum projections. Right: Quantification of the number of puncta in a single focal plane per cell. Error bars represent standard deviation for n > 22 cells quantified. (B) Fluorescence microscopy of cells expressing GFP-tagged endogenous Age2, Gcs1, or Glo3 after treatment with either MNTC or DMSO (control) for 2 minutes. Single focal plane shown. (C) Left: Fluorescence microscopy of Neon-3xFLAG tagged constructs in which the GAP domain of Age2 was swapped with that of Gcs1. Maximum projections. Right: Quantification of the number of puncta in a single focal plane per cell. Error bars represent standard deviation for n > 22 cells quantified. (D) Complementation assay with Neon-3xFLAG-tagged constructs in which the GAP domain of Age2 was swapped with that of Gcs1. All Scale bars, 2 µm. n.s., not significant; *, P ≤ 0.05, **, P ≤ 0.01, ***, P < 0.001; ****, P < 0.0001.

To further explore the mislocalization phenotype of GAP-dead Age2, we tested whether active Arf1 was required for Age2 localization. When we depleted active Arf1 through treatment with the Arf-GEF inhibitor MNTC (Lee et al., 2014; Thomas & Fromme, 2016), we observed that Age2, but not Glo3 or Gcs1, was largely mislocalized to the cytosol (Figure 4B). This suggests Age2 has a unique dependency on Arf1 for its localization.

This led us to ask if Age2 simply requires any functional Arf-GAP domain for its localization. We therefore swapped the GAP domain of Age2 with that of Gcs1 (Figure 4C) and saw partial rescue of Age2 localization. This chimeric construct also rescued growth in the complementation assay, indicating that the partial localization of this chimera is sufficient to provide essential Age2 function in this strain (Figure 4D). Overall, these results indicate that the presence of a functional GAP domain is essential for Age2 localization and function, and that there is some unique aspect of the Age2 GAP domain required for wild-type localization.

### Age2 does not require clathrin or the TGN clathrin adaptors for localization

Age2 is known to function in clathrin-mediated trafficking and may also directly interact with clathrin (Natsume et al., 2006; Poon et al., 2001). We therefore asked whether clathrin plays a role in the recruitment of Age2 to the Golgi. As described above, we found that a loss of the CB had only a minor effect on the punctate localization of Age2 and had no impact on the essential function of Age2 in *gcs1Δ* cells (Figure 2D). To directly test a role for clathrin, we examined Age2 localization in *chc1Δ* cells, which lack the clathrin heavy chain. Although the mutant cells exhibited disrupted Golgi morphology, Age2 remained punctate (Figure 5A). This indicates that Age2 does not require clathrin for localization to Golgi puncta.

**FIGURE 5.**
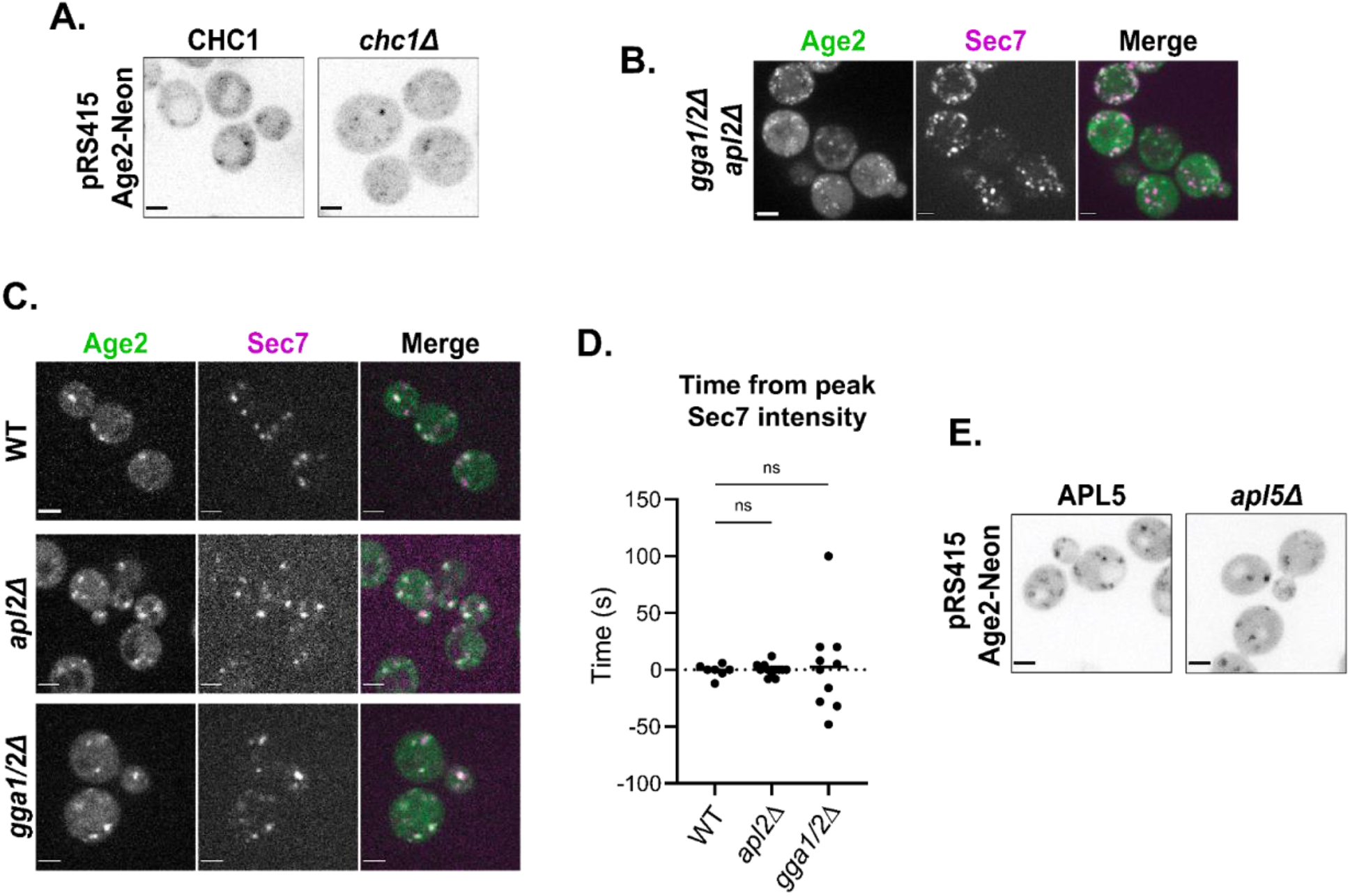
Age2 localization does not require clathrin or clathrin adaptors. (A) Fluorescence microscopy of Age2-Neon expressed via centromeric plasmid in CHC1 or *chc1Δ* cells. Maximum projections. (B) Fluorescence microscopy of Age2-Neon in *gga1Δ gga2Δ apl2Δ* cells. Maximum projections. (C) Fluorescence microscopy of Age2-Neon in *gga1Δ gga2* or *apl2Δ* cells. Single focal plane shown. (D) Peak-to-peak times for Age2 vs Sec7 in WT, *gga1Δ gga2Δ*, or *apl2Δ* cells. Age2 is tagged with Neon in *gga1Δ gga2Δ* and *apl2Δ* strains, and GFP in WT strain. Sec7 is tagged with 6xDsRed in wild-type and *gga1Δ gga2Δ* strains, and RFP-Mars in *apl2Δ* strain. (E) Fluorescence microscopy of Age2-Neon expressed via centromeric plasmid in APL5 or *apl5Δ* cells. Maximum projections. All scale bars, 2 µm.

Other potential candidates for recruiting Age2 to the TGN are the adaptor proteins for the clathrin-mediated trafficking pathways that Age2 is thought to be involved in (Poon et al., 2001). However when both clathrin adaptors at the late-Golgi, GGA and AP-1 (Casler et al., 2019; Casler & Glick, 2020; Daboussi et al., 2012; Day et al., 2018; Highland & Fromme, 2021; Tojima et al., 2019), were absent (in *gga1Δ gga2Δ apl2Δ* cells), Age2 again remained punctate, regardless of the severe disruption in Golgi morphology and cellular function (Figure 5B). We wondered whether clathrin adaptors, while not needed for the punctate localization of Age2, were instead required to confine Age2 to the late-Golgi. We used time-lapse microscopy to measure the dynamics of Age2 throughout Golgi maturation in strains lacking either GGA (*gga1Δ gga2Δ* cells) or AP-1 (*apl2Δ* cells). The timing of Age2 localization was not affected by these mutations, and thus the clathrin adaptors at the late-Golgi do not appear required to recruit Age2 (Figure 5C and D).

Age2 has also been reported to interact with AP-3 (Schoppe et al., 2020), which localizes to the medial- to late-Golgi (Highland & Fromme, 2021). We therefore tested localization of Age2 in a strain lacking AP-3 (*apl5Δ* cells), and Age2 retained punctate localization (Figure 5E).

Taken together, these results indicate that Age2 localization does not require interaction with clathrin or its adaptors, but instead requires the ability to inactivate Arf1 and an interaction between the KKSILSLY amphipathic helix and the membrane surface.

### The levels of active Arf1 rise throughout Golgi maturation

Based on our analysis of the dynamics of Glo3, Gcs1, and Age2, we now have a comprehensive timeline for when the Arf-GAPs localize to the Golgi throughout the maturation process. Earlier work has established the timing of the Arf-GEFs (Gustafson & Fromme, 2017; Highland & Fromme, 2021; Losev et al., 2006). However, we still do not have a good understanding of Arf1 dynamics during Golgi maturation, which in principle is due to the balance of GEF and GAP activity. The major barrier to determining Arf1 localization is that we currently have no way to accurately visualize functional Arf1 in live cells. This is due to the inability to tag Arf1 on either its N- or C-terminus without affecting its function in yeast. The N-terminus of Arf1 is modified with a myristoyl group, which is essential for Arf1 membrane localization and function (Gillingham & Munro, 2007). Fusion tags on the C-terminus of Arf1 have been shown to impair Arf1 function (Hall et al., 2008; Jian et al., 2010). Therefore, in order to visualize Arf1 during Golgi maturation in yeast, a different approach is needed.

As Arf1 is only stably membrane-bound in its active, GTP-bound form, the localization of Arf1 to Golgi compartments is assumed to represent active Arf1-GTP (Antonny, Beraud-Dufour, et al., 1997; Randazzo et al., 1995). To gain an approximate sense of Arf1 dynamics, we used time-lapse microscopy to visualize the dynamics of Arf2 tagged with GFP. Arf2 contributes only ∼10% of the total Arf1/2 pool (Stearns et al., 1990) and thus in contrast to tagging Arf1, tagging Arf2 does not impact cellular function. Arf2-GFP peaked approximately 10 seconds before Sec7, around the medial-to late-Golgi transition (Figure 6, A and E; Supplemental Figure S5A). The caveat to this observation is that Arf2-GFP itself is likely not fully functional.

**FIGURE 6.**
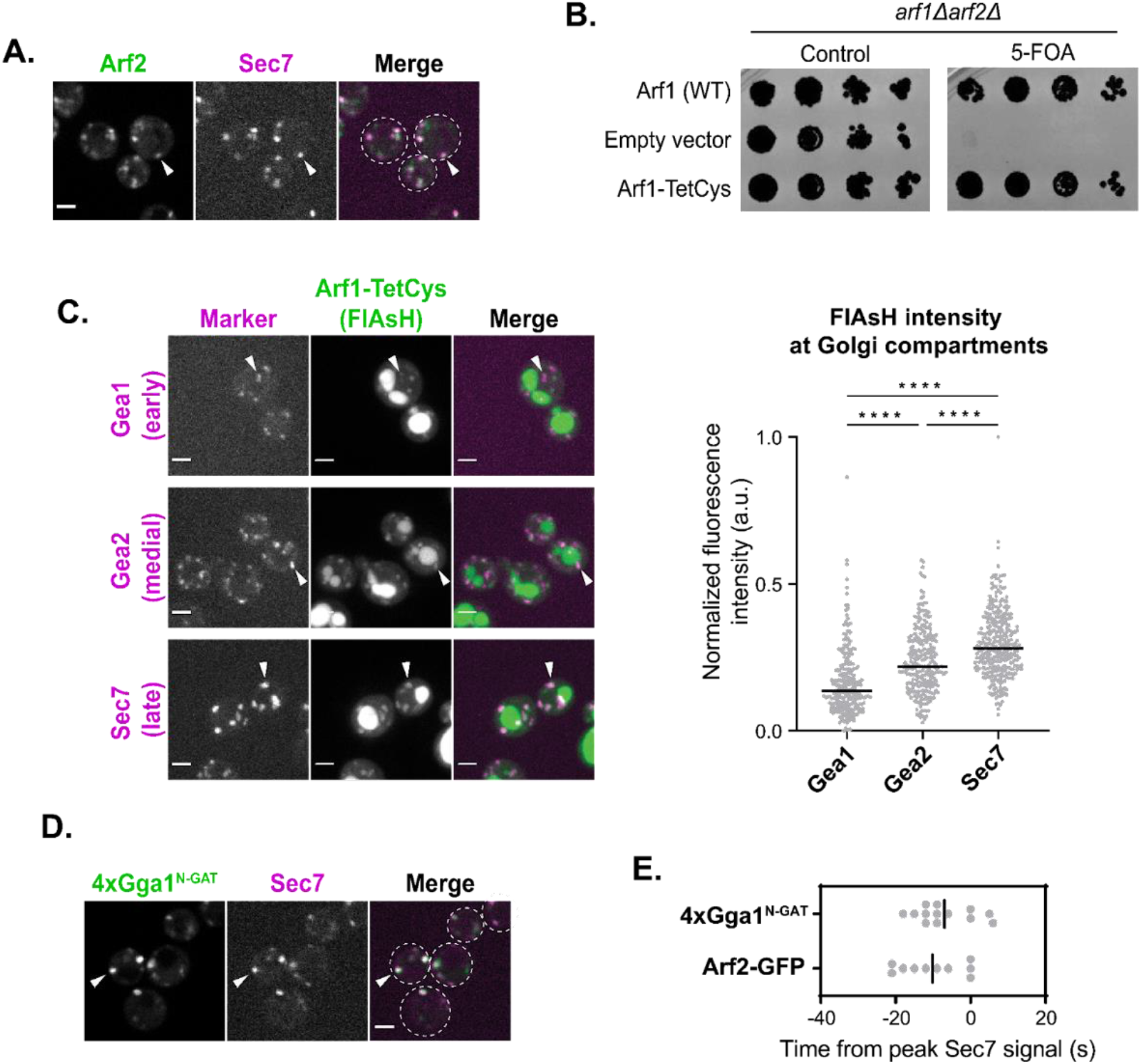
A majority of active Arf1 is at the late-Golgi. (A) Subcellular localization of Arf2-GFP relative to the late-Golgi marker Sec7-6xDsRed. Arrowheads indicate colocalization between tagged proteins. Single focal plane. (B) Complementation assay with Arf1 tagged with TetCys motif for FlAsH labeling. (C) Left: Representative images of FlAsH-labeled cells with Arf1-TetCys alongside markers of the early-(Gea1-3xMars), medial-(Gea2-3xMars), or late-Golgi (Sec7-6xDsRed). Arrowheads indicate colocalization between tagged proteins. Maximum projections. Right: Quantification of intensity of FlAsH-labeling at Gea1-, Gea2-, or Sec7-marked compartments. (D) Subcellular localization of 4xGga1^N-GAT^-Neon relative to late-Golgi marker, Sec7-6xDsRed. Arrowheads indicate colocalization between tagged proteins. (E) Peak-to-peak times for Arf2-GFP and 4xGga1^N-GAT^-Neon versus Sec7-6xDsRed. Line represents the mean. For all images, scale bars, 2 µm. n.s., not significant; *, P ≤ 0.05, **, P ≤ 0.01, ***, P < 0.001; ****, P < 0.0001.

To gain a better understanding of where functional Arf1 localizes, we utilized the FlAsH tag (Griffin et al., 1998). The FlAsH tag is a small fluorescence molecule that binds short tetracysteine (TetCys) motifs that consist of CCXXCC, where X represents any amino acid. We first determined that the fusion of this small TetCys sequence on the C-terminus of Arf1 did not compromise Arf1 function as demonstrated by normal growth in a complementation assay, in contrast to a C-terminal GFP tag which totally abolished growth (Figure 6B; Supplemental Figure S5B). We then examined FlAsH-EDT2 treated cells expressing Arf1-TetCys alongside markers of the early-, medial-, and late-Golgi. FlAsH was found to label Golgi compartments, although there was also nonspecific labeling of the vacuole that was also observed in cells without TetCys-tagged Arf1 (Supplemental Figure S5C). We attempted time-lapse analysis of Arf1-TetCys, but unfortunately the FlAsH signal bleached too quickly to be used for such an analysis. To instead assess the distribution of Arf1 across the Golgi at different stages of maturation, we measured the steady-state fluorescence intensity of FlAsH labeling at each compartment using colocalization with established marker proteins. Arf1 intensity was found to be higher at later compartments relative to earlier compartments (Figure 6C). These results indicate that active Arf1 at the Golgi increases throughout maturation and is consistent with previous static colocalization results (Day et al., 2018).

To assess the dynamics of functional Arf1 localization, we developed an active Arf1-binding probe that consists of the N-terminal portion of the GAT domain of yeast Gga1 (residues 185-230). The corresponding region of human Gga1 was demonstrated to bind Arf1-GTP *in vitro*, and a crystal structure was determined for the Arf1-GTP-GAT domain complex (Collins et al., 2003; T. Shiba et al., 2003). We fused four copies of the GAT domain to mNeonGreen (4xGga1^N-GAT^-Neon) and performed live-cell imaging. We observed that the probe localized to punctate Golgi structures (Figure 6D).

This localization was dependent on Arf1, as the probe was quickly redistributed to the cytosol with MNTC-induced depletion of active Arf1 (Supplemental Figure S5D). Using the ‘Anchor-Away’ method (Haruki et al., 2008), we observed that artificial anchoring of an extra copy of GTP-locked Arf1(Q71L) to the plasma membrane induced the concomitant redistribution of the probe to the PM (Supplemental Figure S5E). Furthermore, we found that the probe could bind directly to Arf1-GTP *in vitro* using a liposome flotation assay with Arf1-loaded liposomes (Supplemental Figure S5F). Arf1 is therefore both necessary and sufficient for localization of this probe *in vitro* and *in vivo*.

Having validated the probe, we proceeded with its use in time-lapse analysis. Similarly to Arf2-GFP, the Arf1-GTP probe peaked at the Golgi ∼7 seconds before Sec7, near the medial- to late-Golgi transition (Figure 6E; Supplemental Figure S6A). We observed that the probe became undetectable during the final stages of Golgi maturation, before the Sec7 signal disappears (Supplemental Figure S6A). This was surprising to us as we expect that Arf1 would still be present during these final stages of Golgi maturation. Our interpretation is that this result may reflect competition between the probe and the endogenous effectors of Arf1-GTP.

Although each approach that we utilized for monitoring the temporal localization of Arf1-GTP has its own caveats, overall, the results of these different approaches for Arf1 visualization are largely in agreement with each other. Taken together our observations indicate that active Arf1 increases throughout Golgi maturation, reaching its highest levels at the medial- to late-Golgi transition (Figure 7).

**FIGURE 7.**
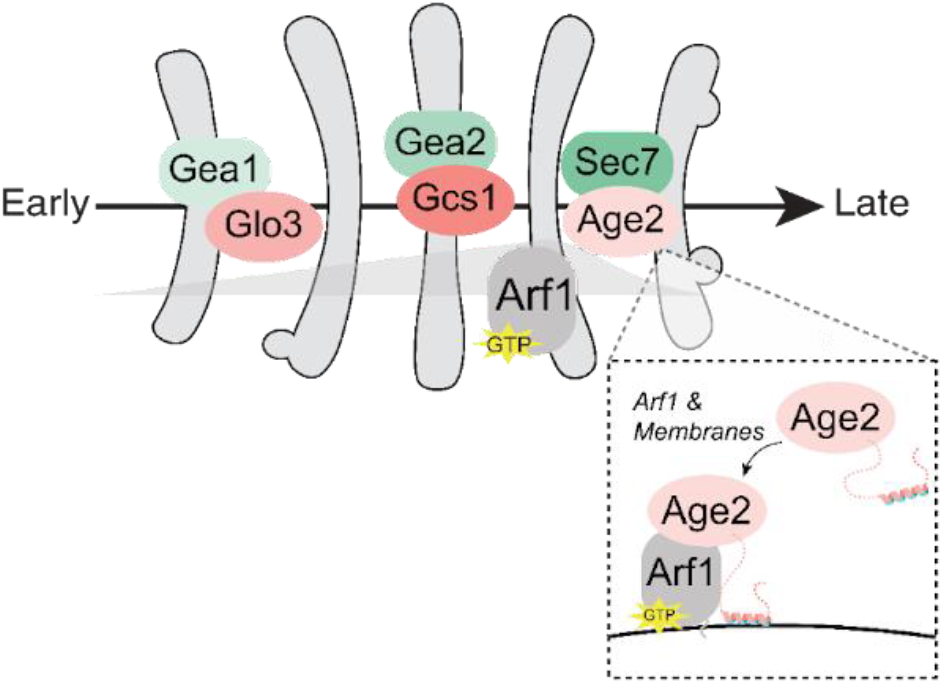
Model for the regulation of Arf1 at the Golgi. As with the GEFs, a specific GAP is present to regulate Arf1 activity at each stage of Golgi maturation. Glo3, Gcs1, and Age2 are mainly localized to the early-, medial-, and late-Golgi, respectively. Age2 localizes to the late-Golgi via interactions with the membrane and Arf1. The resulting activated Arf1-GTP is broadly localized throughout Golgi maturation, but peaks at the medial- to late-Golgi transition. This generally coincides with the expression levels and localizations of the GEFs and GAPs (darker green/pink indicate higher expression levels).

## Discussion

Arf1 is a key regulator of Golgi trafficking and is responsible for initiating several trafficking pathways at distinct times throughout Golgi maturation (Bhave et al., 2014). The work of multiple labs has established the dynamics of the Arf-GEFs as well as several Arf effectors throughout maturation (Casler et al., 2019; Casler & Glick, 2020; Daboussi et al., 2012; Day et al., 2018; Highland & Fromme, 2021; Tojima et al., 2019). However, the maturation dynamics of the Arf-GAPs and of Arf1 itself remained a gap in our understanding of Golgi trafficking. In this study, we address these unknowns to provide a comprehensive view of regulation of Arf1 throughout Golgi maturation.

We determined that a specific Arf-GAP appears responsible for regulating Arf1 at each of the major stages of Golgi maturation, with Glo3 mainly localizing at the early-Golgi, Gcs1 at the medial-Golgi, and Age2 at the late-Golgi. These findings are consistent with previous studies and provide new temporal resolution in the context of the Golgi maturation process.

A caveat of our analysis is that we necessarily focused on the localization of the Arf-GAPs to Golgi compartments, as it is not feasible to quantify the temporal dynamics of fluorescent proteins on fast-moving transport vesicles. However, there are good reasons to assume that Arf-GAP localization on Golgi compartments is directly related to their contributions to trafficking pathways. For example, the COPII coat proteins Sec23 and Sec31, which together act as the GAP for the Arf-family member Sar1, exhibit a steady-state localization on the endoplasmic reticulum at ER-exit sites that produce COPII vesicles (Kurokawa et al., 2016; Nishikawa & Nakano, 1991; Rossanese et al., 1999). Similarly, Arf-GAPs appear to be incorporated into the COPI coat during vesicle biogenesis (Arakel et al., 2019; Dodonova et al., 2017; Spang et al., 2010; Weimer et al., 2008). Therefore, although Arf-GAPs are expected to localize to COPI and clathrin vesicles, they clearly localize strongly to the Golgi compartments which give rise to these vesicles. Our observation that the Arf-GAPs appeared as multiple small puncta per Golgi compartment is also consistent with their localization representing their incorporation into budding vesicles, as these small discrete puncta resemble sites of vesicle formation. In addition to ensuring their incorporation into nascent vesicles, the Golgi localization of Arf-GAPs may also serve to counterbalance GEF activity (Nie & Randazzo, 2006).

The unique localizations of each of the GAPs are undoubtedly key aspects of their function. The localization of Gcs1 and Glo3 have been attributed to interactions with curved membranes and COPI, respectively (Weimer et al., 2008; Xu et al., 2013). However, the preference of Gcs1 for curved membranes does not fully explain how Gcs1 is restricted to predominantly medial Golgi compartments, and its localization is likely to be a result of multiple contributing interactions. For example, Gcs1 localization is also influenced by a positively charged residue upstream of the curvature sensing ALPS motif, suggesting specificity for anionic membrane surfaces (Xu et al., 2013).

Most studies on the Golgi Arf-GAPs have centered on ArfGAP1/Gcs1 and ArfGAP2/3/Glo3, and thus our knowledge of the Age2 localization mechanism has remained limited. The human homolog SMAP2 has been reported to localize to the TGN and early endosomes (Funaki et al., 2011; Natsume et al., 2006). Yeast has a minimal endomembrane system in which the TGN serves as the early endosome (Day et al., 2018), and therefore Age2 and SMAP2 localize to functionally equivalent compartments. Golgi localization of SMAP2 was proposed to be achieved through a protein-protein interaction between a basic stretch within the GAP domain (KYEKKK) and an unidentified TGN protein (Funaki et al., 2011). However, AlphaFold predicts several of these residues to be important for structural integrity of the GAP domain (Jumper et al., 2021; Varadi et al., 2021) and mutating these residues therefore likely disrupts protein folding.

We have now determined the functional regions of Age2 that are essential for targeting the late-Golgi. We determined that GAP activity and a conserved “KKSILSLY” sequence are each essential for the Golgi localization of Age2. Although a previous report found that this sequence was not required for localization of SMAP2 (Funaki et al., 2011), this prior study used overexpression of SMAP2 constructs in cell culture. We found that this sequence appears to fold into an amphipathic helix that mediates the direct binding of Age2 to the Golgi membrane. Interestingly, the helix is relatively short compared to other known membrane-binding amphipathic helices, and alteration of its primary sequence that preserved its amphipathicity abolished its ability to target Age2 to the Golgi. This observation may suggest the existence of an additional binding partner for the KKSILSLY sequence, or a strict requirement of the specific physical properties of the helix for membrane insertion.

We were surprised to observe that Age2 localization is dependent on its GAP activity. In principle, a GAP-dead mutant would be expected to still localize properly and bind to Arf1 but be incapable of carrying out catalysis (Arakel et al., 2019). As GAP activity is required for release of a vesicle coat and subsequent fusion, one possibility is that the apparent cytosolic localization of the GAP-dead mutant could be explained by its entrapment on coated vesicles, which cannot be resolved by fluorescence microscopy due to their small size and rapid diffusion. This possibility is supported by the identification of Age2 on AP-3 vesicles and an increase in AP-3 vesicles with the loss of Age2 (Schoppe et al., 2020). It is possible that the GAP domain may also participate in other interactions that contribute to Age2 localization. The inability of the GAP domain of Gcs1, whose localization was not sensitive to MNTC, to fully rescue Age2 localization suggests that either the Age2 GAP domain interacts more tightly with Arf1, or that one or both of the GAP domains form additional interactions with other proteins.

In order to determine the temporal dynamics of Arf1 throughout maturation, we turned to the small FlAsH tag to minimize the invasiveness of tagging Arf1 (Griffin et al., 1998). FlAsH labeling of Arf1 tagged with TetCys revealed that the amount of active Arf1 at the Golgi increases at later Golgi compartments. The TGN serves as the site of several Arf-dependent outgoing trafficking pathways, and thus an increase in Arf1 may be required to support the high trafficking demand.

Rapid photobleaching of FlAsH precluded its use for time-lapse analyses of Arf1 maturation dynamics. To circumvent this issue, we developed a probe for Arf1-GTP based on the Arf1-binding portion of Gga1. Assessment of the maturation dynamics of the Arf probe corroborated the finding that more Arf1 is present later in maturation and further revealed that Arf1 peaked in localization at the medial-to-late-Golgi. Surprisingly, the Arf probe became undetectable at the Golgi before Sec7, the late-Golgi Arf-GEF. We expected that active Arf1 would be present at the Golgi whenever an activating Arf-GEF was. This observation may indicate that the probe is outcompeted by other Arf1 effectors, and thus implies that the highest levels of freely available active Arf1 are found at the medial- to late-Golgi. This suggests that Arf1 is quickly consumed by vesiculation at the end of the lifetime of the compartment and is consistent with the known timing of several Arf1 dependent trafficking pathways which occur at the late-Golgi/TGN (Daboussi et al., 2012; Highland & Fromme, 2021; Tojima et al., 2019).

The mechanism by which Arf1 recruits the different effectors required for these pathways to the Golgi at precise and distinct times is not known for all effectors. In general, it is thought that coincidence detection is the main mechanism for this specificity (D’Souza-Schorey & Chavrier, 2006). We speculate that the availability of Arf1 at the Golgi may also contribute, and competition between effectors for free Arf1 may help to refine their localizations. Additionally, it is essential that there is sufficient Arf1 to facilitate all Arf1-initiated pathways. A simple model is that the amount of active Arf1 at the Golgi is a product of its activation by GEFs and inactivation by GAPs. Our now complete picture of the localization of the Arf-GEFs, Arf-GAPs, and resulting Arf, provides support for this general model (Figure 7). The order of the Arf-GEFs from low to high expression levels - Gea1, Gea2, and Sec7 - corresponds to their localization at the early-, medial-, and late-Golgi respectively (Bui et al., 2009; Gustafson & Fromme, 2017; Ho et al., 2018). Consistently, we found that the product of the Arf-GEFs, active Arf1, also increases later in maturation. In contrast, the late-Golgi Arf-GAP Age2 has the lowest expression levels, followed by the early Arf-GAP Glo3, and the medial-Arf-GAP Gcs1 has the highest expression (Ho et al., 2018).

This work provides a detailed picture of the spatiotemporal regulation of Arf1 in budding yeast. Overall, the localization of the Arf-GEFs and Arf-GAPs dictate when and how much active Arf1 is localized to different compartments. While these GEFs and GAPs each share their highly conserved catalytic domains, they possess several other unique domains that determine their localization. A common feature of the Arf-GEFs and Arf-GAPs is their direct interaction with membranes, often through amphipathic helices (Bigay et al., 2005; Drin et al., 2007; Kliouchnikov et al., 2009; Muccini et al., 2022). Arf-GEFs also rely on interactions as effectors of other GTPases (Christis & Munro, 2012; Cohen et al., 2007; Gustafson & Fromme, 2017; Hofmann et al., 2007; Li et al., 2007; McDonold & Fromme, 2014), while GAPs appear to more often interact with vesicle coat components (Eugster et al., 2000; Inoue & Randazzo, 2007; Natsume et al., 2006; Watson et al., 2004). Overall, both Arf-GEFs and Arf-GAPs generally possess multiple interactions that are important for their proper targeting, and future studies are needed to fully characterize their localization mechanisms.

## Materials and Methods

### Yeast strains and plasmids

All yeast strains and plasmids (listed in Supplemental Tables S1 and S2, respectively) were constructed using conventional methods. All strains are in the SEY6210 (MATα) or SEY6210.1 (MATa) background, with a few exceptions noted here: The wild-type, *chc1*Δ, and *apl5*Δ strains used in Figures 5A and 5E were in a BY4741a background. To purify Age2, Age2 (WT or mutant) expression plasmids were transformed into BJ5459, a protease deficient strain (Hickey et al., 2009; Jones, 1991). Lastly, for the anchor-away experiment, a rapamycin-insensitive yeast strain expressing Pma1-2xFKBP was used (Auffarth et al., 2014; Haruki et al., 2008).

Yeast strains were constructed by mating or genetic manipulation via homologous recombination. All *S. cerevisiae* fluorescent fusion proteins were expressed under the endogenous promoter created by C-terminally tagging the endogenous gene via homologous recombination of PCR-amplified pFA6 cassettes (Longtine et al., 1998). An exception is the Age2-Neon-3xFLAG mutants, which were instead integrated as a single copy via integration plasmids into strains lacking endogenous Age2 and were expressed under the native AGE2 promoter with an ADH1 terminator. The Arf1 probe (4xGga1N-GAT-Neon) was also integrated via an integration plasmid and was expressed under the YPT1 promoter and terminator. The functionality of all proteins were verified by normal cell growth and/or protein localization.

### Microscopy

Yeast were grown to log phase in synthetic media at 30°C. Cells were imaged at log phase at room temperature. Images were acquired with a CSU-X spinning-disk confocal system (Intelligent Imaging Innovations) with a DMI6000 B microscope (Leica Microsystems), 100× 1.46 NA oil immersion objective, and an Evolve 512 Delta EMCCD camera (Photometrics). Images were captured using Slidebook 6 software (Intelligent Imaging Innovations). For time-lapse series, single focal planes were captured every 1 - 4 seconds for a total of 3 - 5 minutes.

For MNTC experiments, cells were treated with 20 µM 6-methyl-5-nitro-2-(trifluoromethyl)-4H-chromen-4-one (MNTC) (MolPort-000-729-160; dissolved in dimethyl sulfoxide [DMSO]) or just DMSO as a control. Cells were imaged in a glass-bottomed dish, and MNTC or DMSO was added directly to the dish (time = 0).

For FlAsH labeling experiments, cells were grown to log phase overnight in synthetic media containing 4 μM FlAsH-EDT2 (Cayman Chemical cat #20704) dissolved in DMSO. During all steps preceding imaging, cells were kept in the dark to prevent bleaching of FlAsH-EDT2.

Before imaging, cells were washed with fresh synthetic media and allowed to incubate, shaking for 10 minutes, twice. Cells were finally resuspended in fresh synthetic media and imaged.

For anchor-away experiments, FRB-Mars-ΔN13Arf1(Q71L) was relocalized to the PM by rapamycin induced dimerization with Pma1-2xFKBP12. Cells were treated with 1 μg/mL rapamycin for 45 min before imaging.

In general, maximum projections (6 focal planes, 0.4 µm apart) are shown for images in which the degree of “punctateness” is being assessed, and single focal planes are shown for images in which the colocalization between two proteins is being assessed.

### Image analysis

Time-lapse analysis was performed using Slidebook 6 software (Intelligent Imaging Innovations). Golgi compartments were deemed suitable for analysis if they remained within the focal plane for the entirety of the compartment’s lifetime and did not overlap with other Golgi compartments. A region of interest (ROI) was created to capture the compartment, and the fluorescence intensity over time was measured for each channel. For each channel, the intensity values were normalized from 0 to 1. The peak-to-peak times were measured as the time between the maximum intensities in each channel.

Quantification of the number of puncta per cell in an individual focal plane was done using ImageJ (FIJI). For each image, the focal plane in which the cells were in clearest focus was selected. Trainable Weke segmentation was used to identify objects deemed as puncta. To train a classifier model, pixels in an image of cells expressing WT Age2-Neon-3xFLAG cells were selected and categorized as either “puncta’’ or “background”. This classifier model was then applied to images of each strain that was analyzed. A consistent threshold was then applied on the resulting probability maps to create a mask of all segmented objects. Objects of fewer than 4 pixels were removed from the analysis. For each cell, the number of objects (puncta) were counted. If a daughter cell was more than half the size of the mother, they were treated as two separate cells. At least 22 cells were counted for each analyzed strain.

Assessment of intensity of FlAsH-labeled Arf1-TetCys was performed using Slidebook 6. For each compartment marker (Gea1, Gea2, or Sec7), a threshold was manually set to create a mask that selected a majority of the marked Golgi compartments without also including cytosolic signal. For each compartment marker, the mean intensity of FlAsH-labeling was measured. As FlAsH-EDT2 also accumulated in the vacuole, marked compartments that overlapped with vacuolar signal were excluded to avoid artificial measurements. To do this, a threshold was manually set to include the relatively bright vacuolar signal. As other FlAsH labeled structures were also selected, objects were required to be at least 1 μM in size to include only vacuoles. Finally, the measured intensity values were background subtracted by subtracting the mean intensity of a ROI that did not include any cells. Background subtracted intensity values were normalized from 0 to 1.

### Yeast complementation assays

For complementation (“plasmid shuffling”) assays with Age2 mutants, integration plasmids harboring mutant versions of Age2 were integrated into a sensitized background (*age2*::HIS3, *gcs1*::KANMX, [pRS416 *AGE2*] (pCF1066)) and grown on -LEU plates. Neither *AGE2* or *GCS1* is essential, but the loss of both genes is synthetically lethal, and thus functional Age2 is required for growth in this background. Growth is maintained by the presence of pRS416 *AGE2*. For each strain, two clones each from two independent integrations were tested for growth, and only one representative clone is shown in the figures above. Serial dilutions of each strain were replicated on -LEU and 5-FOA, in which the maintenance plasmid is counter-selected against. Cells were grown for 3 days at 30°C.

For complementation assays with tagged Arf1 constructs, centromeric plasmids were transformed into a sensitized background (*arf1*Δ::HIS3, *arf2*Δ::KANMX, Sec7-Mars::TRP1 [pRS416 *ARF1*] (pCF1022)) and grown on -LEU plates. *ARF1* and *ARF2* are an essential paralogous gene pair, with either gene required for growth, and growth of this strain is maintained by pRS416 *ARF1*. For each transformation, several colonies were serially diluted and replicated on -LEU and 5-FOA. The assay was performed with three individual transformations for each construct tested. Cells were grown for 3 days at 30°C.

### Protein purifications

Age2 was purified from yeast expressing GST-Age2 under the GAL1 promoter. Twelve liters of cells were grown in a nutrient rich raffinose containing media (1% yeast extract, 2% peptone, 2% raffinose) at 30°C to OD600 ∼ 0.8 - 1.2. Expression was induced with 0.2% galactose for 5 hours. Cells were harvested by centrifugation, washed, and resuspended in lysis buffer (10% glycerol, 50 mM Tris pH 8, 450 mM NaCl, 1% CHAPS, 20 mM MgCl2, 5 mM ATP, 1 mM Dithiothreitol [DTT], 1X Roche Protease Inhibitor Cocktail, 1 mM 4-(2-aminoethyl)benzenesulfonyl fluoride [AEBSF]). The cell suspension was frozen dropwise in liquid N2 and lysed using a freezer mill (SPEX SamplePrep). The lysate was clarified by centrifugation and incubated with glutathione resin (G-Biosciences cat #786-310) for 1 hour. The resin was washed with lysis buffer. To remove bound chaperones, the resin was washed with 20 resin volumes of room temperature chaperone removal buffer (50 mM Tris pH 7.5, 400 mM NaCl, 0.01% CHAPS, 20 mM MgCl2, 50 mM KCl, 5 mM ATP, 1 mM DTT) three times. Finally, to reduce phosphorylation, the resin was washed with CIP buffer (10% Glycerol, 50 mM Tris pH 8, 450 mM NaCl, 1% CHAPS, 20 mM MgCl2, 1 mM DTT) and incubated with Quick CIP (NEB cat #M0525S) for ∼16 hours. To remove CIP, the resin was washed with Purification Buffer (50 mM Tris pH 7.5, 500 mM NaCl, 0.004% CHAPS, 5% glycerol, 1 mM DTT). Protein was eluted by incubating with PreScission protease for at least 4 hours.

GST-4xGga1N-GAT-Neon was produced in Rosetta 2 cells. One liter of cells were grown in LB-Miller media at 37°C to OD600 ∼ 0.6 - 0.8. Cells were induced with 0.2 mM IPTG at 18°C for ∼16 hours. Cells were then harvested by centrifugation, washed, and resuspended in lysis buffer (1X Phosphate-buffered saline [PBS], 5 mM β-mercaptoethanol [β-ME], 1 mM phenylmethylsulfonyl fluoride [PMSF]). Cells were lysed by sonication and clarified by centrifugation. The cleared lysate was then incubated with glutathione resin (G-Biosciences cat #786-310) for 3 hours. The resin was washed with lysis buffer and then PreScission Cleavage buffer (50 mM Tris-HCl pH 7.5, 150 mM NaCl, 1 mM EDTA, 1mM DTT). Protein was eluted by incubation with PreScission protease for ∼16 hours.

Myristoylated Arf1 and soluble ΔN17-Arf1 were purified using previously established methods (Ha et al., 2005; Richardson et al., 2012; Richardson & Fromme, 2015).

### Liposome preparations

Liposomes were prepared as described previously (Richardson & Fromme, 2015). Lipids and 1% DiR near-infrared dye in chloroform were mixed in the molar ratios described in Supplemental Table S3. Experiments were performed using liposomes prepared from a Folch lipid extract (Sigma Aldrich Cat #B1502-25MG), except for the experiment shown in Supplemental Figure S5F, which instead used a lipid mix designed to mimic the composition of the late-Golgi/TGN (Klemm et al., 2009). The lipid mixes were then vacuum dried and rehydrated in HK (20 mM HEPES, pH 7.4, 150 mM KOAc) buffer overnight at 37°C. Lipids were finally extruded through 100 nm filters and stored at 4°C.

### Tryptophan fluorescence Arf-GAP assay

Arf-GAP assays were performed based on methods described previously (Paczkowski et al., 2012; Richardson & Fromme, 2015): A native tryptophan fluorescence assay (297.5 nm excitation, 340 nm emission) was used to monitor the nucleotide state of Arf1 (Antonny, Huber, et al., 1997). For GAP assays with myristoylated Arf1, myrArf1-GTP was first activated on liposomes by adding HKM buffer (20 mM HEPES, pH 7.4, 150 mM KOAc, 2 mM MgCl2), 200 μM 100 nm Folch liposomes, 2 μM Arf1, 100 μM GTP, and 2 mM EDTA sequentially and incubating at 30°C for 20 minutes. To stop the exchange, 5 mM MgCl2 was added, and the mixture was stored on ice throughout the experiment. The assays were performed at 30°C, and Arf1-loaded liposomes were warmed to 30°C for 5 minutes before being transferred to a cuvette and beginning measurements. Age2 (WT or mutant) was added to 0.2 μM (t = 0). The buffer that the protein is purified in was used instead for mock experiments. Analysis was performed in GraphPad Prism, in which the traces were fitted to a one-phase exponential decay curve to determine the rate of GAP activity.

For GAP assays with ΔN17-Arf1, a truncated form of Arf1 that is unable to bind membranes, no liposomes were required for Arf1 activation. For the ΔN17-Arf1 GAP assays, 1.5 μM Age2 was used, but otherwise these assays were carried out similarly.

### Liposome flotation assays

To assess membrane recruitment of Age2, liposome flotation was used as previously described (Highland et al., 2023; Richardson & Fromme, 2015). For this, 80 μL reactions with HK buffer +1 mM DTT, 250 μM 100 nm Folch liposomes, and 10 μg Age2 were incubated for 20 minutes at 30°C. Liposomes and bound protein were separated from unbound protein via discontinuous sucrose gradient flotation. 2.5 M sucrose in HK was added to the reaction to a final concentration of 1 M (“Input” sample) and transferred to 7 × 20 mm polycarbonate centrifuge tubes (Beckman Coulter cat #343775). HK buffer with 0.75 M sucrose was gently laid on top of the reaction mixture followed by a layer of HK, and centrifuged at 390,000 × g/20°C for 20 min. Lipids were recovered from the top layer (“Membrane-bound” sample). Samples were analyzed via SDS-PAGE, and proteins were visualized by Coomassie and lipids were visualized by DiR dye.

To assess the Arf1-dependent recruitment of 4xGga1N-GAT-Neon to membranes (Supplemental Figure S5F), liposome floatation was carried out similarly but with the following modifications. First, 3 µg myristoylated Arf1 was loaded with the nonhydrolyzable GTP analog GMP-PNP or GDP via EDTA-mediated nucleotide exchange in the presence of TGN liposomes. Then 6 µg of the purified 4xGga1N-GAT-Neon construct was added to the reaction mixture and allowed to incubate as described above. HKM buffer was used instead of HK in the sucrose gradients. Finally, to quantify membrane recruitment, protein levels were normalized to the amount of lipids to account for variations in lipid recovery.

### Western blot analysis of whole cell extracts

To monitor the expression levels of Age2 mutants, 5 ODs of log phase yeast were harvested by centrifugation, washed, and resuspended in 1 mL cold water. To precipitate protein, 110 μL of 100% Trichloroacetic Acid [TCA] was added and the cells were incubated on ice for at least 30 minutes. Precipitated protein was collected and resuspended in cold acetone via sonication. Precipitate was collected again by centrifugation and the supernatant discarded. Pellets were dried using a SpeedVac to remove residual acetone. To resuspend pellets and prepare samples for SDS-PAGE, pellets were vortexed and boiled at 95°C for 5 minutes each with Boiling Buffer (50 mM Tris pH 7.5, 1 mM EDTA, 1% SDS) and 0.5mm glass beads. 2X Urea Buffer (150 mM Tris pH 6.8, 6 M Urea, 6% SDS, Bromophenol Blue) was then added and the sample was vortexed again and finally heated at 65°C for 10 minutes. Samples were analyzed via SDS-PAGE and western blot probing for FLAG epitopes. Consistent loading of protein samples was verified by blotting for Glucose-6-Phosphate Dehydrogenase (G6PDH).

### Antibodies

The 1° mouse anti-FLAG antibody (Sigma Aldrich Cat #F3165-1MG) used to detect Age2-Neon-3xFLAG was used at a 1:2000 dilution. The 1° rabbit anti-G6PDH antibody (Sigma Aldrich Cat #SAB2100871) was used at a concentration of 1:30,000. The 2° sheep IgG horseradish peroxidase-linked anti-mouse antibody was used at a 1:5000 dilution (Amersham cat #NXA931V). The 2° donkey IgG horseradish peroxidase-linked anti-rabbit antibody was used at a 1:5000 dilution (Amersham cat #NA934V).

### Statistical analysis

Statistical tests were performed using GraphPad Prism 8 software. For all figures, ns indicates not significant; *, P < 0.05; **, P < 0.01; ***, P < 0.001; ****, P < 0.0001. Statistical significance was determined using an unpaired t test with Welch’s correction (Figure 2B, Figures 3, C-E, Figure 4A, and Supplemental Figure S5F), or a one-way analysis of variance with Tukey’s test for multiple comparisons (Figure 4C and Figure 6C) or Dunnet’s test for multiple comparisons (Figure 2D, Figure 3A, and Figure 5D)

## Author Contributions

K.M.M. and J.C.F. conceptualized the study. K.M.M. designed and performed the experiments, analyzed and interpreted the results, prepared the figures, and wrote the manuscript with advice from J.C.F., who also secured funding and revised the manuscript.

## Acknowledgements

We thank the laboratories of J. Baskin, A. Bretscher, S. Emr, C. Sevier, and C. Ungermann for sharing advice, equipment, strains, and reagents. We are grateful to Fromme lab members for helpful discussions and for sharing reagents (B. Brownfield prepared myrArf1 and generated CFB3518; A. Allen prepared ΔN17-Arf1). This work was supported by NIH/NIGMS grant R35GM136258 to J.C.F. and a National Science Foundation Graduate Research Fellowship to K.M.M.

## Supplemental Figures

**Supplemental Figure 1.**
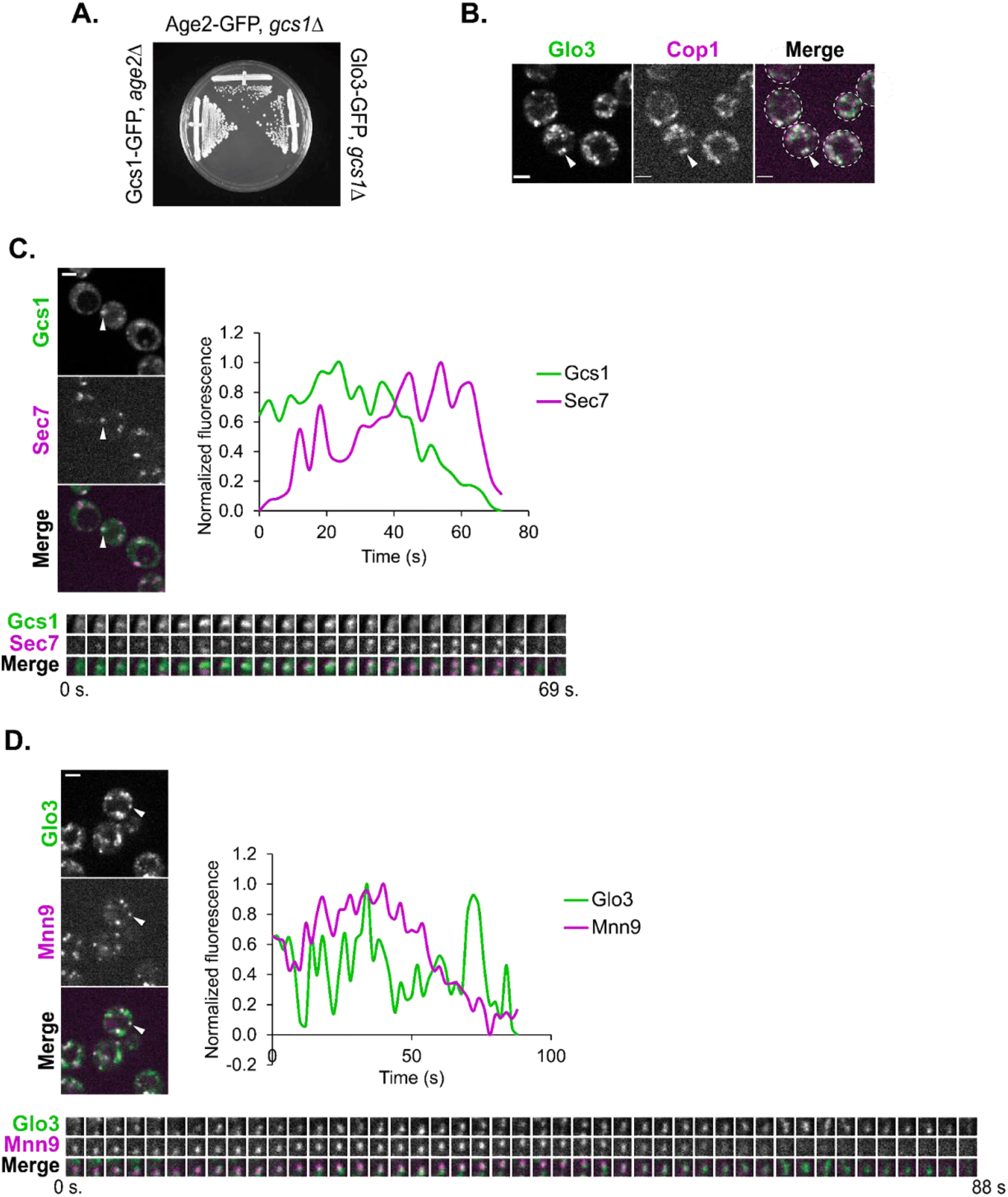
Timelapse analyses of Arf-GAP maturation dynamics at the Golgi. (A) Growth assay of tagged Arf-GAPs in strains lacking another Arf-GAP that together form an essential pair. (B) Subcellular localization of Glo3-Neon relative to the COPI subunit, Cop1-mCherry. (C) Left: Representative image of time-lapse microscopy of Gcs1-GFP versus Sec7-6xDsRed. Arrowhead denotes Golgi compartment of interest. Bottom: imaging of the compartment of interest over time. Right: plot of normalized fluorescence intensity in the compartment of interest over time. (D) Left: Representative image of time-lapse microscopy of Glo3-Neon versus Mnn9-mCherry. Arrowhead denotes Golgi compartment of interest. Bottom: imaging of the compartment of interest over time. Top right: plot of normalized fluorescence intensity in the compartment of interest over time.

**Supplemental Figure 2.**
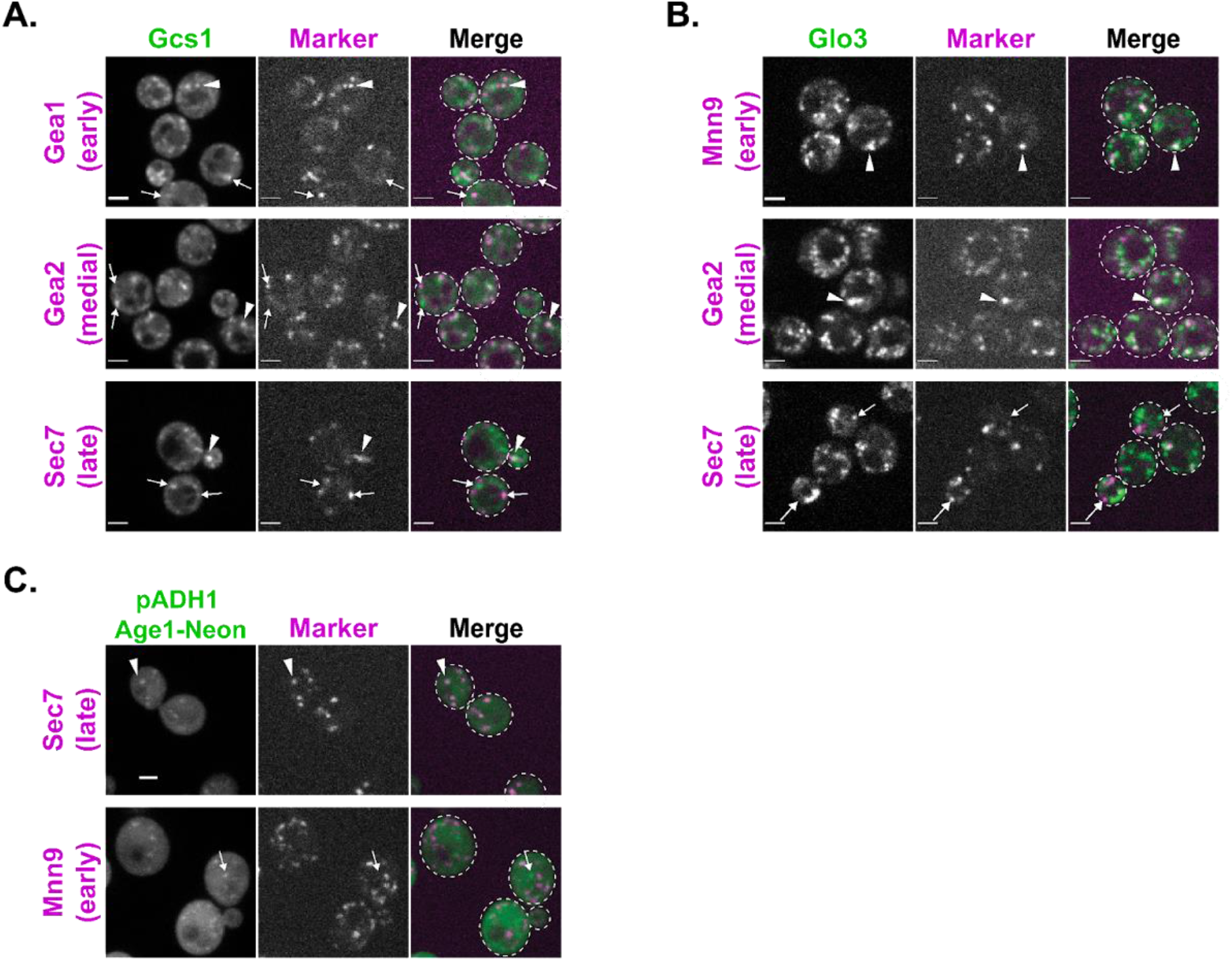
Colocalization analysis of Arf-GAPs and Golgi compartment markers. (A) Subcellular localization of Gcs1 relative to early- (Gea1-3xMars), medial- (Gea2-3xMars), or late- (Sec7- 6xDsRed) Golgi markers. Gcs1 is tagged with GFP in the Gea1-3xMars strain, and Neon in the Gea2-3xMars and Sec7-6xDsRed strain. Arrowheads and arrows indicate colocalization or a lack of colocalization between proteins, respectively. Single focal planes. (B) Subcellular localization of Glo3 relative to early- (Mnn9-mCherry), medial- (Gea2-3xMars), or late- (Sec7-6xDsRed) Golgi markers. Glo3 is tagged with GFP in the Sec7-6xDsRed strain, and Neon in all other strains. Arrowheads and arrows indicate either colocalization or a lack of colocalization between proteins, respectively. Single focal planes. (C) Subcellular localization of Age1 overexpressed on a centromeric plasmid under the strong ADH1 promoter relative to markers of the late- (Sec7-6xDsRed) or early- (Mnn9-mCherry) Golgi. Maximum projections. For all images, scale bars represent 2 µm.

**Supplemental Figure 3.**
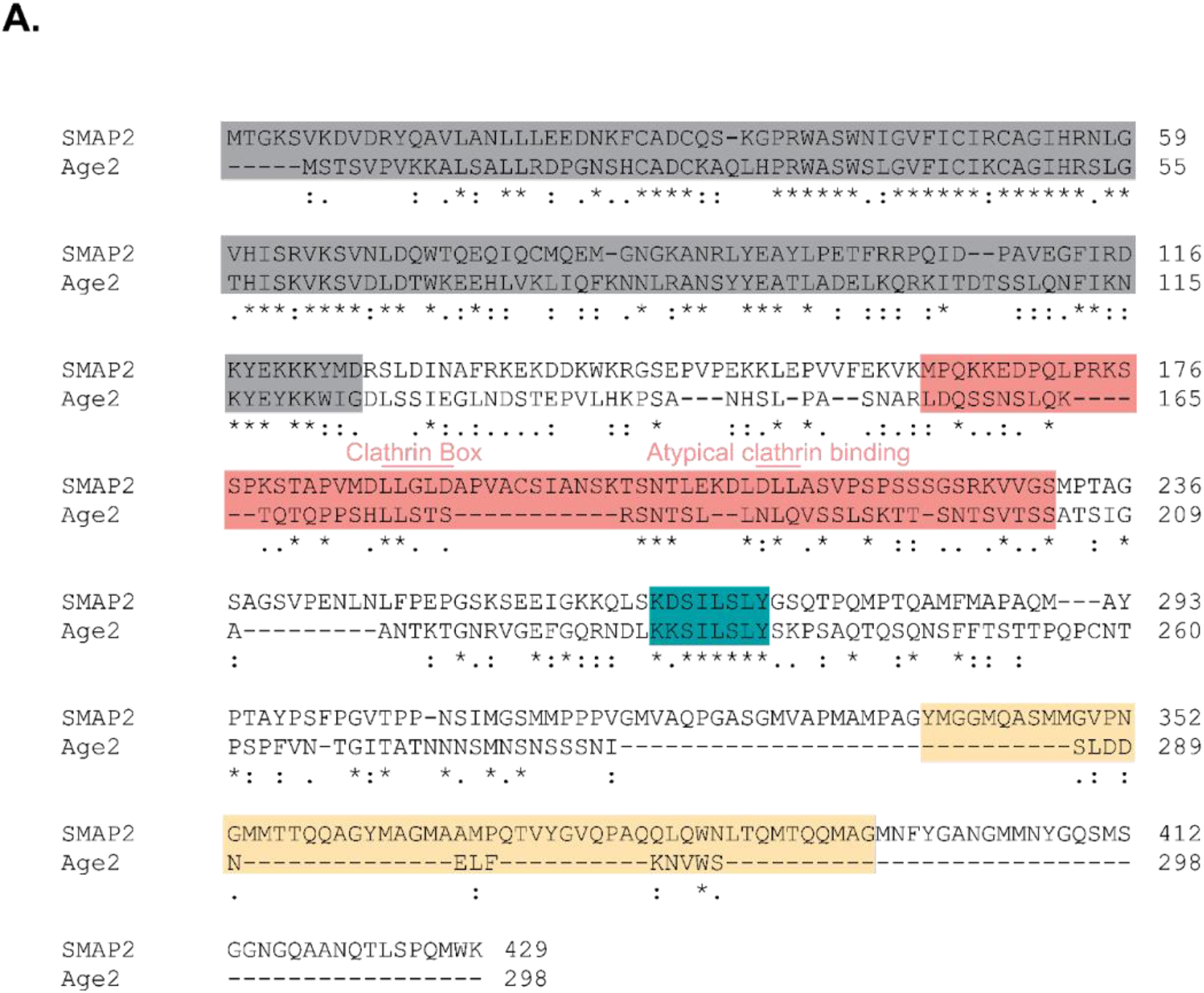
Alignment of Age2 and SMAP2.

(A) Alignment of Age2 and SMAP2 (accession numbers P40529 and Q8WU79 respectively). SMAP2 functional domains (Natsume et al., 2006) are highlighted (GAP domain in gray, clathrin interaction domain in pink, and CALM interaction domain in yellow). The conserved KKSILSLY sequence is also highlighted in teal. The clathrin interacting motifs within the clathrin interaction domain are denoted.

**Supplemental Figure 4.**
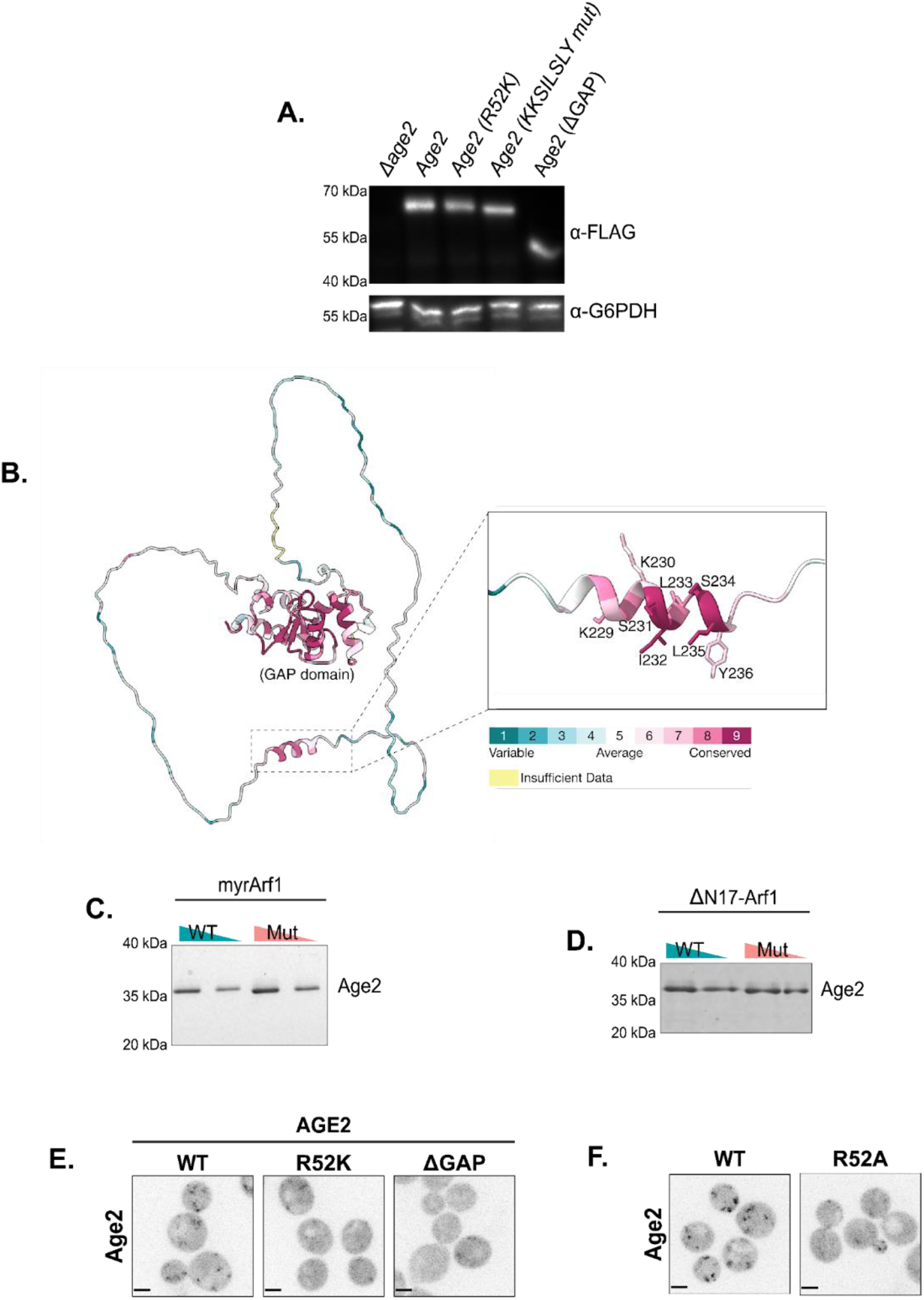
Further validation of the dependency of Age2 on Arf1 and membranes for localization. (A) Western blot analysis of whole cell lysates expressing Age2-Neon-3xFLAG mutants using anti-FLAG. Glucose-6-Phosphate Dehydrogenase (G6PDH) serves as a loading control. (B) AlphaFold prediction of Age2 structure, with a close up of the predicted helix formed by the conserved KKSILSLY sequence (Jumper et al. 2021; Varadi et al. 2021). Model is colored by conservation as determined by ConSurf (Glaser et al., 2003; Landau et al., 2005). (C) SDS-PAGE of purified proteins used for GAP assays with myrArf1. Triangles indicate the relative amount of protein loaded. (D) SDS-PAGE of purified protein used for GAP assays with ΔN17-Arf1. Triangles indicate the relative amount of protein loaded. (E) Fluorescence microscopy of Age2-Neon-3xFLAG mutants expressed as an extra copy in an otherwise wild-type *AGE2* strain. Maximum projections. (F) Fluorescence microscopy of GAP dead Age2(R52A)-Neon-3xFLAG. Maximum projections. For all images, scale bars represent 2 µm.

**Supplemental Figure 5.**
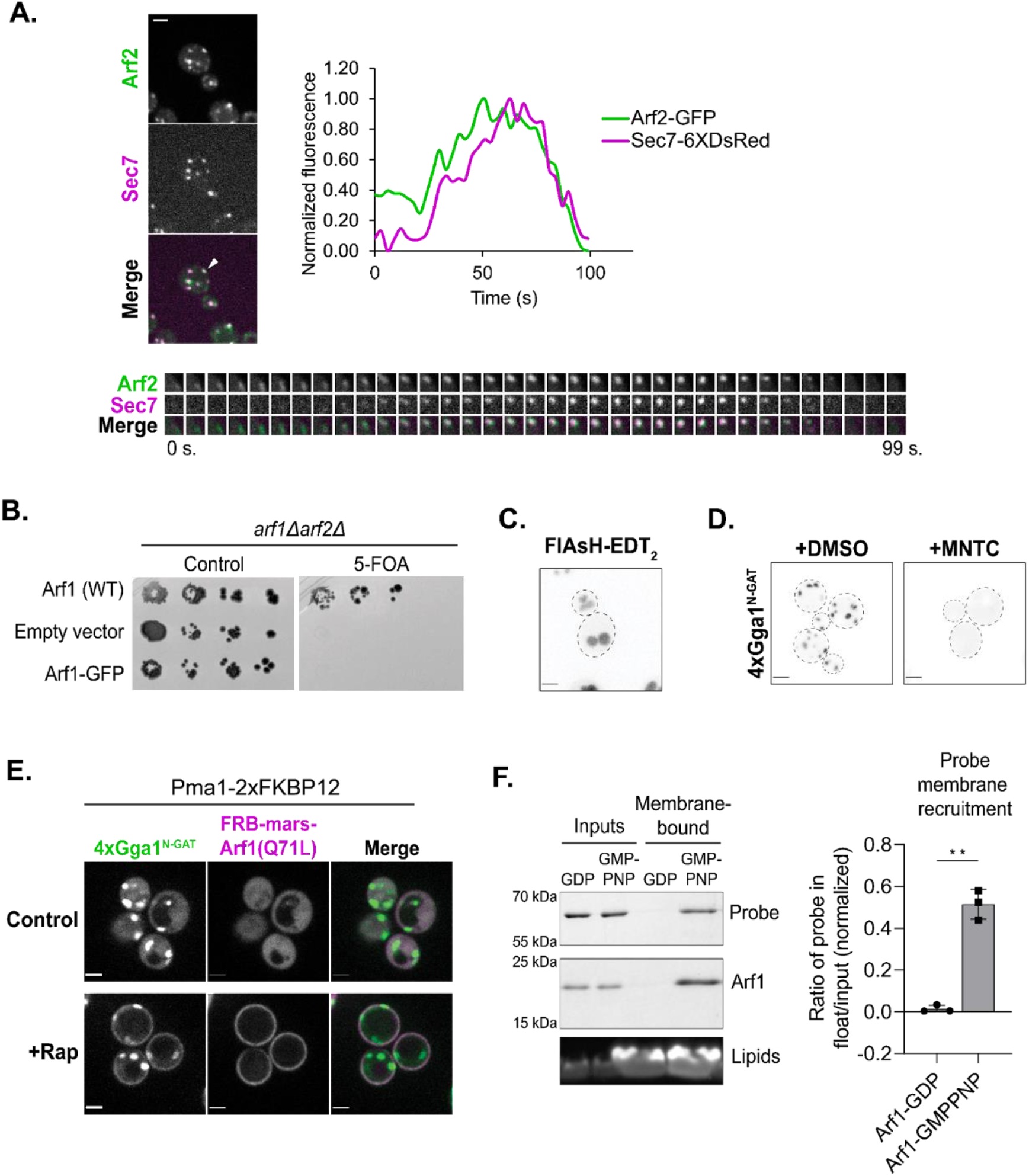
Validation of FlAsH-labeling approach and 4xGga1^N-GAT^-Neon probe. (A) Left: Representative image of time-lapse microscopy of Arf2-GFP versus Sec7-6xDsRed. Arrowhead denotes Golgi compartment of interest. Bottom: Imaging of the compartment of interest over time. Top right: Plot of normalized fluorescence intensity in the compartment of interest over time. (B) Complementation assay with Arf1 tagged with GFP. (C) Representative image of FlAsH-treated wild-type cells showing vacuolar staining. Maximum projection. (D) Fluorescence microscopy of 4xGga1^N-GAT^-Neon after treatment with either MNTC or DMSO (control) for 1 minute. (E) Representative images of 4xGga1^N-GAT^-Neon in cells expressing FKBP-tagged Pma1 and FRB-tagged GTP-locked (Q71L) Arf1 treated with rapamycin to induce dimerization of FKBP and FRB and thus the relocalization of Arf1 to the plasma membrane. Single focal plane. (F) Left: *In vitro* liposome flotation assay to assess binding of purified protein to membranes in an Arf1-dependent manner. Right: Quantification of 4xGga1^N-GAT^-Neon probe membrane recruitment, as measured by coomassie band intensity ratio. Error bars represent standard deviation for n = 3 assays.

**Supplemental Figure 6.**
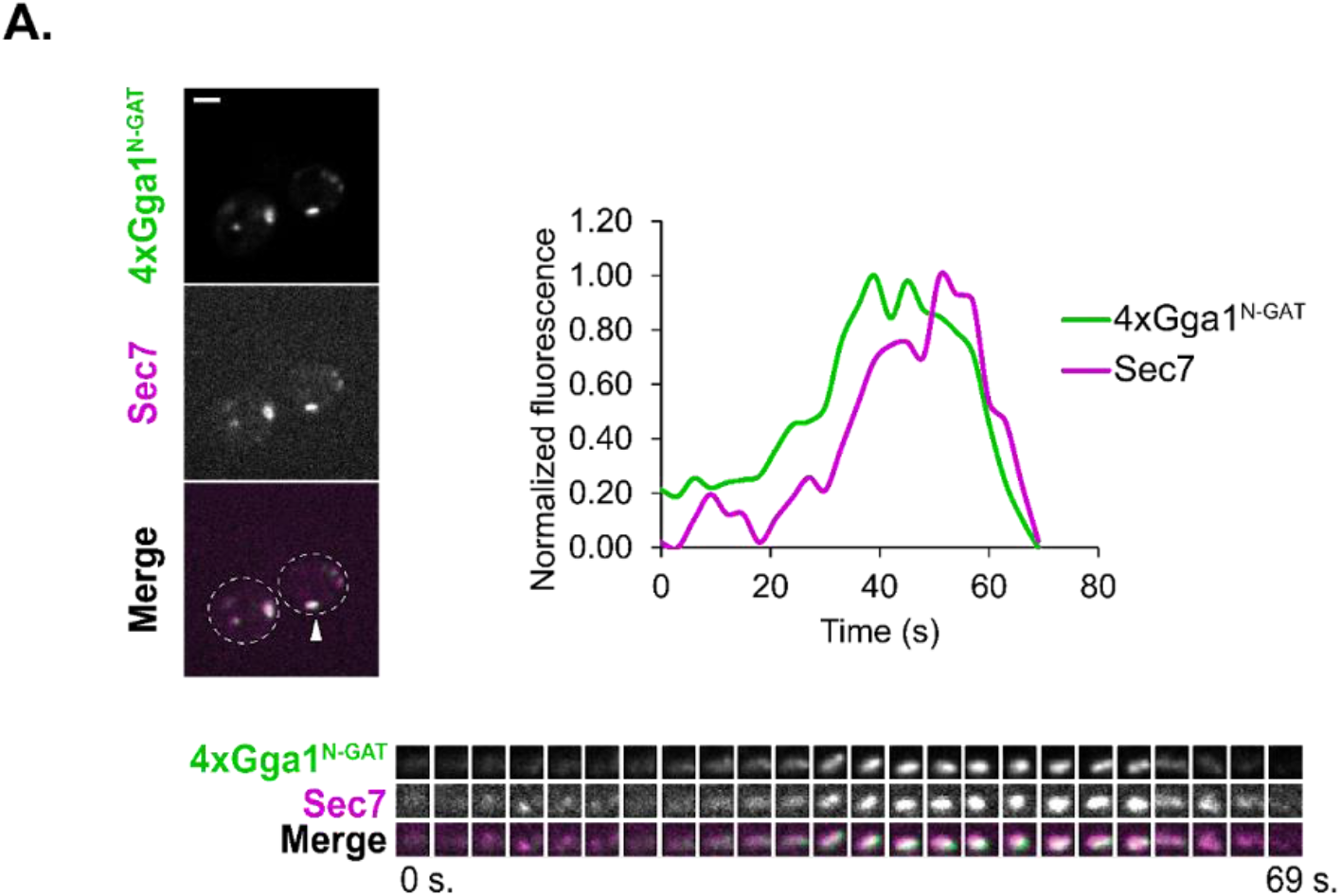
Time-lapse analysis of 4xGga1^N-GAT^ maturation dynamics. (A) Left: Representative image of time-lapse microscopy of 4xGga1^N-GAT^-Neon versus Sec7-6xDsRed. Arrowhead denotes Golgi compartment of interest. Bottom: Imaging of the compartment of interest over time. Top right: Plot of normalized fluorescence intensity in the compartment of interest over time.

**Table S1.**
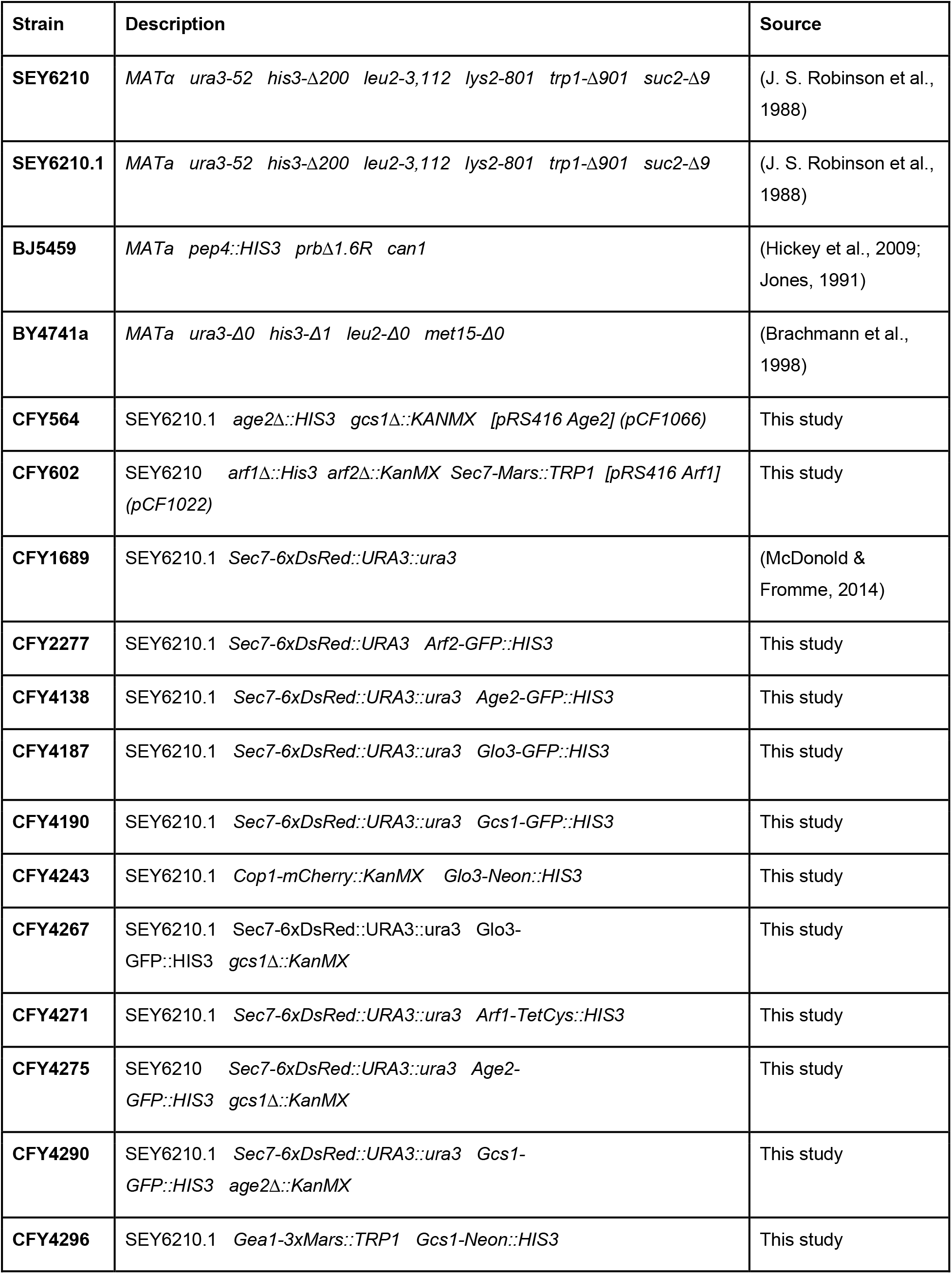
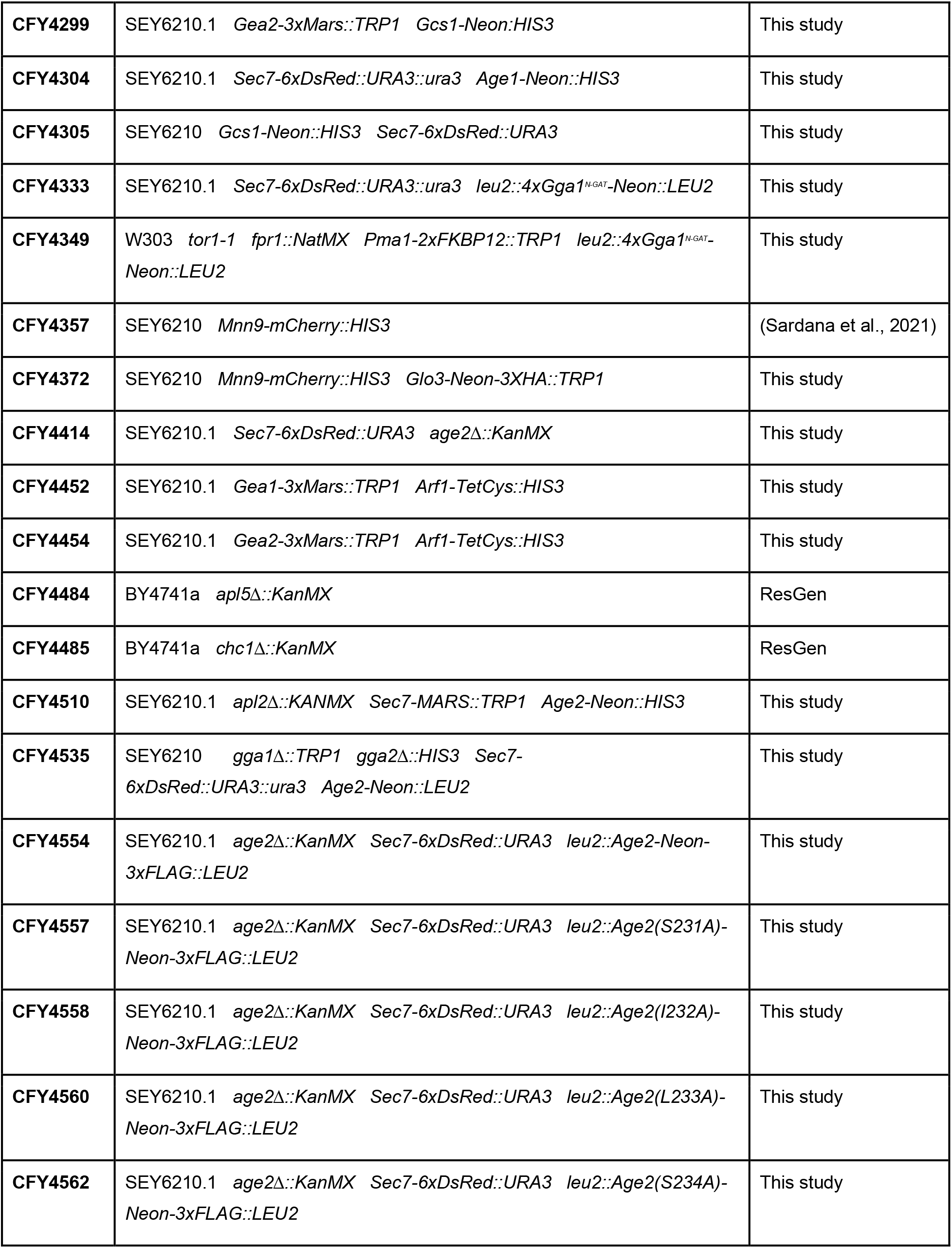
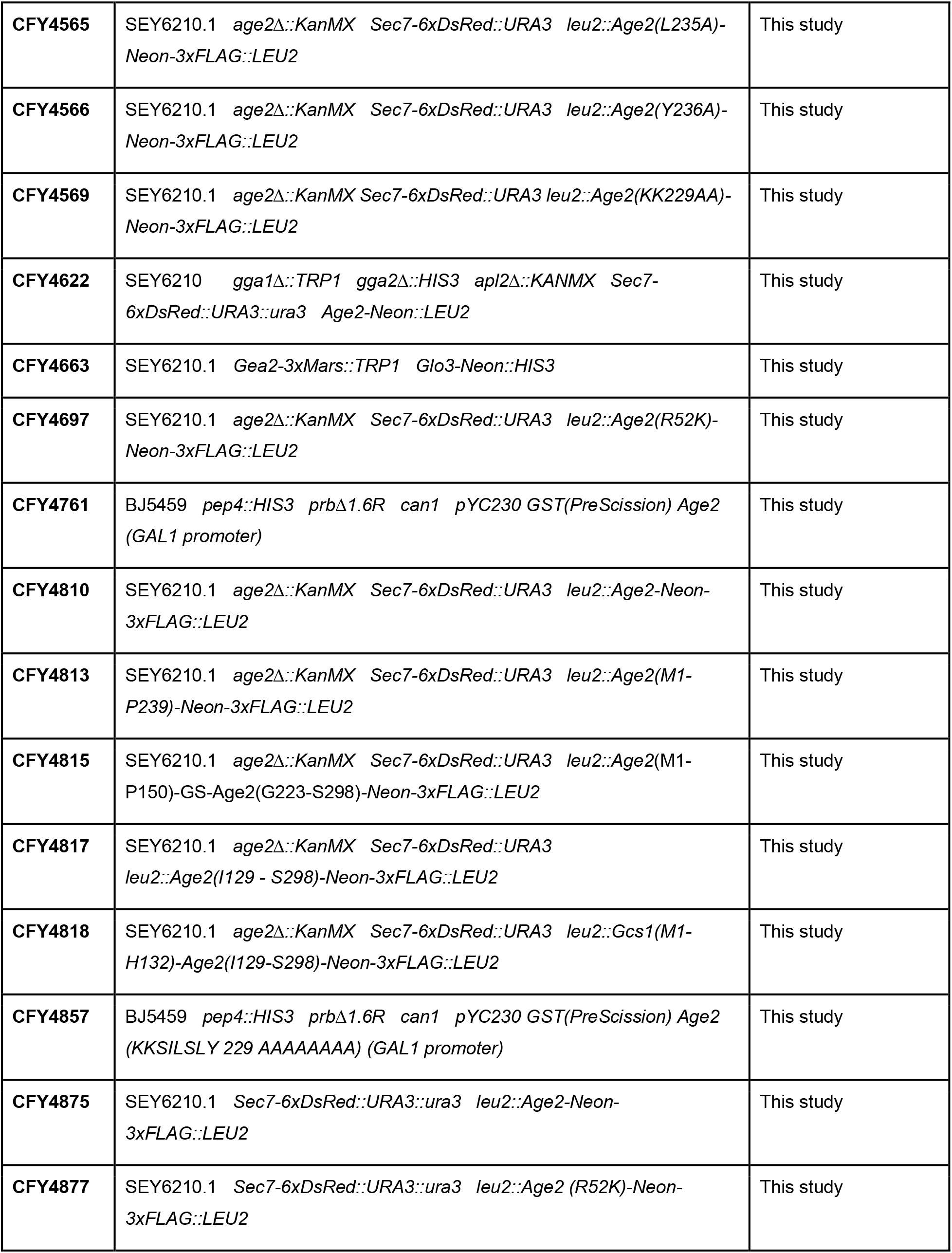
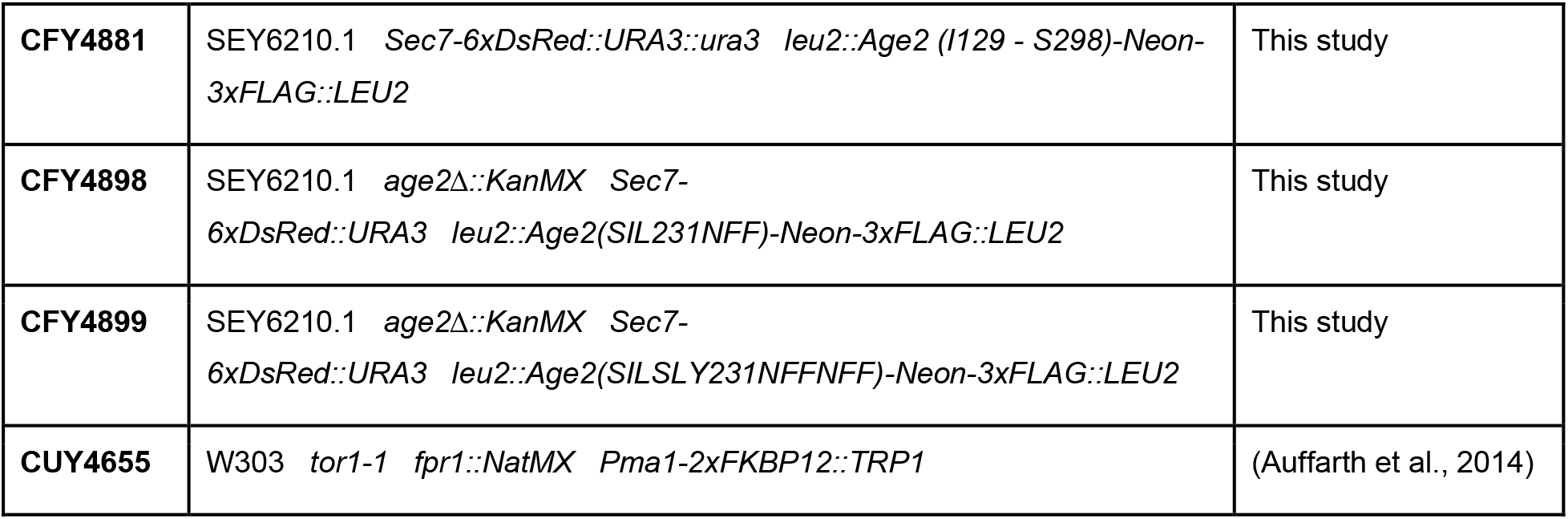
Yeast strains used in this study:

**Table S2.** Plasmids used in this study:

| Name | Description | Vector Backbone | Source |
| --- | --- | --- | --- |
| **pRS305** | Integration vector (LEU2) | pRS305 | (Sikorski & Hieter, 1989) |
| **pRS414** | Vector (TRP1) | pRS414 | (Sikorski & Hieter, 1989) |
| **pRS415** | Vector (LEU2) | pRS415 | (Sikorski & Hieter, 1989) |
| **pArf1** | Arf1 (Expression plasmid) | pET3 | (Weiss et al., 1989) |
| **pNmt1** | Nmt1 (Expression plasmid) | pCYC | (Duronio et al., 1990) |
| **pLT1** | FRB-Mars-Arf1(Q71L)ΔN13 (GPD promoter, CYC1 terminator) | pRS423 | (Thomas et al., 2021) |
| **CFB3518** | Arf1-GFP | pRS415 | B. Brownfield |
| **pCF1022** | Arf1 | pRS416 | (Thomas & Fromme, 2016) |
| **pCF1023** | Arf1 | pRS415 | (Thomas et al., 2021) |
| **pCF1053** | ΔN17-Arf1 (Expression plasmid) | pET28a | (Richardson et al., 2012; Richardson & Fromme, 2015) |
| **pCF1066** | Age2 | pRS416 | This study |
| **pKM38** | Arf1-GSSG-TetCys | pRS415 | This study |
| **pKM55** | 4xGga1<sup>N-GAT</sup>-Neon (Integration plasmid) | pRS305 | This study |
| **pKM73** | 4xGga1<sup>N-GAT</sup>-Neon (Expression plasmid) | pGEX-6P-1 | This study |
| **pKM90** | Age2-Neon (ADH1 terminator) | pRS415 | This study |
| **pKM92** | Age2-Neon-3xFLAG (Integration plasmid) | pRS305 | This study |
| **pKM93** | Age2 (S231A)-Neon-3xFLAG (Integration plasmid) | pRS305 | This study |
| **pKM94** | Age2 (I232A)-Neon-3xFLAG (Integration plasmid) | pRS305 | This study |
| **pKM95** | Age2 (L233A)-Neon-3xFLAG (Integration plasmid) | pRS305 | This study |
| **pKM96** | Age2 (S234A)-Neon-3xFLAG (Integration plasmid) | pRS305 | This study |
| **pKM97** | Age2 (L235A)-Neon-3xFLAG (Integration plasmid) | pRS305 | This study |
| **pKM98** | Age2 (Y236A)-Neon-3xFLAG (Integration plasmid) | pRS305 | This study |
| **pKM99** | Age2 (KK229AA)-Neon-3xFLAG (Integration plasmid) | pRS305 | This study |
| **pKM127** | Age2 (R52K)-Neon-3xFLAG (Integration plasmid) | pRS305 | This study |
| **pKM131** | Age2 (M1-P239)-Neon-3xFLAG (Integration plasmid) | pRS305 | This study |
| **pKM132** | Age2(M1-P150)-GS-Age2(G223-S298)-Neon-3xFLAG (Integration plasmid) | pRS305 | This study |
| **pKM133** | Age2 (KKSILSLY 229 AAAAAAAA)-Neon-3xFLAG (Integration plasmid) | pRS305 | This study |
| **pKM135** | GST [PreScission] Age2 (GAL1 promoter) | pYC230 | This study |
| **pKM136** | Age2 (I129 - S298)-Neon-3xFLAG (Integration plasmid) | pRS305 | This study |
| **pKM145** | Age2(M1-P150)-GS(72aa)-Age2(G223-S298)-Neon-3xFLAG (Integration plasmid) | pRS305 | This study |
| **pKM146** | Age1-Neon (ADH1 promoter, CYC1 terminator) | pRS414 | This study |
| **pKM152** | GST [PreScission] Age2 (KKSILSLY 229 AAAAAAAA) (GAL1 promoter) | pYC230 | This study |
| **pKM154** | Age2 AH (RNDLKKSILSLYSK) (2X)-Neon (YPT1 promoter and terminator) | pRS415 | This study |
| **pKM156** | Age2 (SIL231NFF)-Neon-3xFLAG (Integration plasmid) | pRS305 | This study |
| **pKM157** | Age2 (SILSLY231NFFNFF)-Neon-3xFLAG (Integration plasmid) | pRS305 | This study |

*All plasmids (besides plasmids for protein expression) express genes under their endogenous promoter and terminator unless otherwise denoted.*

**Table S3.** Compositions of liposomes used in this study.

| Lipid | TGN | Folch |
| --- | --- | --- |
| DOPC | 24 | - |
| POPC | 6 | - |
| DOPE | 7 | - |
| POPE | 3 | - |
| DOPS | 1 | - |
| POPS | 2 | - |
| DOPA | 1 | - |
| POPA | 2 | - |
| PI | 29 | - |
| PI(4)P | 1 | - |
| CDP-DAG | 2 | - |
| PO-DAG | 4 | - |
| DO-DAG | 2 | - |
| Ceramide (C18) | 5 | - |
| Cholesterol | 10 | - |
| Folch Lipids (Folch fraction I) | - | 99 |
| DiR Dye | 1 | 1 |

